# Targeting the conserved active site of splicing machines with specific and selective small molecule modulators

**DOI:** 10.1101/2023.06.21.545906

**Authors:** Ilaria Silvestri, Jacopo Manigrasso, Alessandro Andreani, Nicoletta Brindani, Marco De Vivo, Marco Marcia

## Abstract

The self-splicing group II introns are bacterial and organellar ancestors of the nuclear spliceosome and retro-transposable elements of pharmacological and biotechnological importance. Integrating enzymatic, crystallographic, and simulation studies, we demonstrate how these introns recognize small molecules through their conserved active site. These RNA-binding small molecules selectively inhibit the two steps of splicing by adopting distinctive poses at different stages of catalysis, and by preventing crucial active site conformational changes that are essential for splicing progression. Our data exemplify the enormous power of RNA binders to mechanistically probe vital cellular pathways. Most importantly, by proving that the evolutionarily-conserved RNA core of splicing machines can recognize small molecules specifically, our work puts solid bases for the rational design of splicing modulators not only against bacterial and organellar introns, but also against the human spliceosome, which is a validated drug target for the treatment of congenital diseases and cancers.

## INTRODUCTION

Splicing is a ubiquitous and essential biological reaction whereby protein-coding or regulatory RNAs (the exons) are excised from precursor transcripts by removing intronic sequences. In prokaryotes and eukaryotic organelles, splicing is performed by auto-catalytic introns, some of which regulate the expression of vital metabolic genes (Chillón I., 2021). Of these, the so-called group II introns constitute the ancestors of the eukaryotic spliceosome, a megadalton-large ribonucleoprotein complex that ensures the correct maturation of ∼90% of human genes and whose aberrant activity causes ∼15% of all human hereditary diseases and cancers (Jiang and Chen, 2021). Besides catalyzing their self-splicing, group II introns also act as mobile retroelements, thus crucially contributing to genetic diversity throughout evolution (Martinez-Abarca and Toro, 2000). Importantly, as splicing ribozymes and retroelements, group II introns are potential targets of antifungal agents and molecular machines that can be engineered for site-specific insertion of cargo genes into genomic DNA (Chillón I., 2021). Therefore, their modulation with small RNA-targeting molecules has enormous potential to probe fundamental biological mechanisms, with direct translational applications in biotechnology and human medicine (Warner et al., 2018).

Importantly, small organic compounds have recently been shown to target nuclear and organellar splicing complexes (Fedorova et al., 2018; Naryshkin et al., 2014). Such breakthroughs led to the approval of the drug risdiplam for the treatment of neuromuscular disorders, and to the design of the lead compound intronistat B for inhibiting the growth of pathogenic fungi (Fedorova et al., 2018; Naryshkin et al., 2014). However, the mode of action and mechanism of these few compounds remain uncertain.

As a matter of fact, the exact comprehension of whether small molecules can specifically and/or selectively bind RNA remains quite challenging (Falese et al., 2021; Manigrasso et al., 2021b). RNA-binding molecules typically tend to be unspecific and promiscuous, by interacting in a sequence-independent manner with the phosphodiester backbone of RNA or by intercalating stacked nucleobases inducing structural misfolding (Warner et al., 2018). Moreover, describing the principles of RNA recognition by small organic compounds and elucidating their mode of action at the molecular level is severely impaired by the availability of only 746 high-resolution structures of RNA-ligand complexes in the Protein Data Bank (PDB) as of April 2^nd^, 2023.

In this context, we and others have previously demonstrated that both the eukaryotic spliceosome and the bacterial group II introns possess an evolutionarily and structurally-conserved active site (Marcia et al., 2021; Marcia and Pyle, 2012, 2014). This RNA catalytic core binds a cluster of di- and mono-valent metal ions, which acts as an indispensable cofactor for splicing (Marcia et al., 2021; Marcia and Pyle, 2012, 2014). Both splicing machines follow the same reaction chemistry, which consists of two steps. During the first step of splicing, the 5’-exon is excised through a nucleophilic reaction that induces a transiently-inactive ‘toggled’ state, necessary to recruit the reactants of the second step (Manigrasso et al., 2020; Marcia and Pyle, 2012; Marcia et al., 2013b). Then, in the second step of splicing, the free 5′-exon attacks the 3′-splice junction, producing ligated exons and a free intron (Chillón I., 2021). This vital and multistep process involves sequence-specific interactions within a highly-structured metal-aided catalytic site, which in principle offers unique opportunities of favorable RNA-ligand interactions.

Here, we now demonstrate experimentally that the conserved RNA-based active site of splicing ribozymes can indeed recognize small organic compounds specifically and selectively. These compounds anchor to catalytic nucleotides and metal ions, compete with the splice junctions, and selectively prevent active site structural rearrangements required for splicing progression. Our work thus reveals an unexpected and unprecedented mode of action of splicing molecular modulators, through their direct targeting of the splicing site. These unforeseen findings, obtained through the integration of enzymatic, computational, and high-resolution crystallographic studies, determine the exact mechanism of inhibition of bacterial and organellar splicing, with important implications for the design of antifungal antibiotics. More broadly, they also provide the molecular bases for future design of spliceosomal active site modulators that establish nucleotide-specific interactions around the splice junctions. Such targeting approach, which was impossible to practically envision until now, has great potential for the treatment of congenital neurological disorders and cancer that derive from alternative splicing defects.

## RESULTS

### Bacterial and organellar splicing is inhibited by small molecules in a step-specific manner

Following the recent identification of small molecule inhibitors of mitochondrial group IIB introns (Fedorova et al., 2018), here we set out to elucidate their molecular mechanism and to explore whether these compounds could be used as novel molecular probes to gain mechanistic insights into the different steps of the splicing reaction, an approach that had been possible until now only through conventional mutagenesis of active site residues or through replacement of active site metal ions (Marcia et al., 2021; Marcia and Pyle, 2012, 2014).

We first tested whether these molecules inhibit bacterial group IIC introns, besides their mitochondrial homologues, because the I1 intron from *Oceanobacillus (O.) iheyensis* can be readily crystallized and would thus enable us to obtain mechanistic insights at high-resolution. Using an established radio-analytic self-splicing assay, we determined that the most active compound, i.e. intronistat B, inhibits *O. iheyensis* group IIC intron with a *K*_i_ = 1.700 ± 0.004 μM (**Figures 1 and S1**, **Table S1**, see chemical structure of intronistat B in **Figure 5**). For reference, intronistat B inhibits the ai5γ group IIB intron from *Saccharomyces (S.) cerevisiae* with *K*_i_ = 0.360 ± 0.020 μM (Fedorova et al., 2018). Of note, an orthogonal FRET assay, which probes the multiple-turnover spliced-exon reopening (SER) reaction also catalyzed by group II introns, allowed us to confirm an IC_50_ value of 2.5 ± 0.7 μM for the ai5γ group IIB intron, similar to previous reports (Fedorova et al., 2018). Unfortunately, this assay did not work for the *O. iheyensis* group IIC intron, possibly owing to the fact that the shorter exon-binding site sequence (EBS1) in this intron makes the SER reaction less efficient (Popovic and Greenbaum, 2014). We have thus estimated an apparent IC_50_ of 2.3 ± 0.4 μM for the *O. iheyensis* group IIC intron, as calculated from the radio-analytic self-splicing assay at 15 minutes post-addition of the compound. Importantly, we further established that intronistat B targets the folded intron in its active state. This is demonstrated by its inhibitory effect, which is evident also when the compound is added to an ongoing splicing reaction (**Figure S2**).

**Figure 1.**
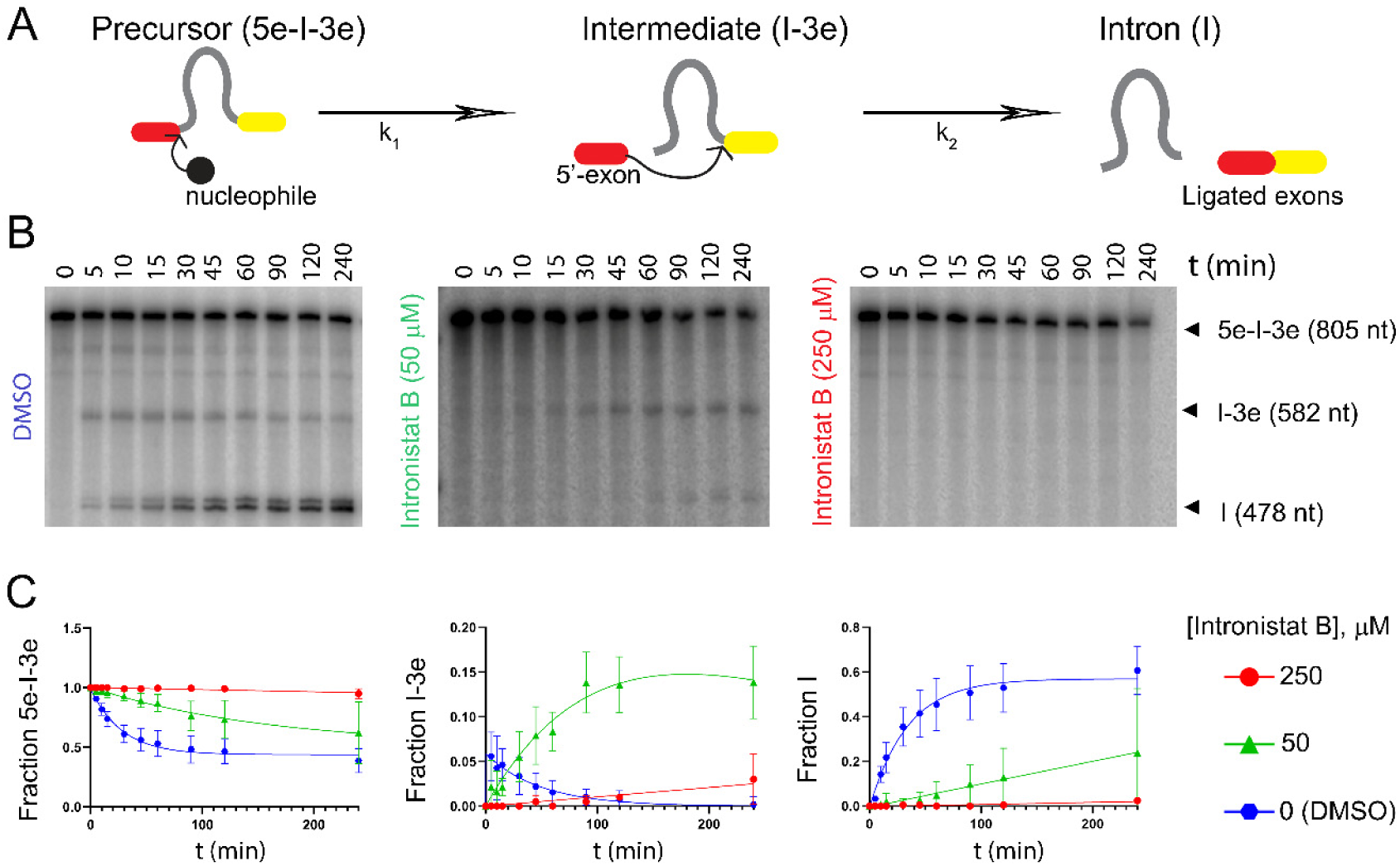
Kinetics of splicing inhibition. **(A)** Schematic of the splicing reaction. k_1_ is the rate constant of the first and k_2_ of the second step of splicing. Kinetic rate constants relevant to our study are reported in **Table S1**. **(B)** Representative splicing kinetics in the absence (control, DMSO) and in the presence of intronistat B (50 and 250 μM). Precursors are indicated as 5e-I-3e (nt length in parenthesis). Intermediate (I-3e) and linear intron (I) migrate as double bands because of cryptic cleavage sites, as explained previously (Manigrasso et al., 2020; Marcia and Pyle, 2012). **(C)** Evolution of the populations of precursor (5e-I-3e, left panel), intermediate (I-3e, middle panel), and linear intron (I, right panel) over time. Error bars represent standard errors of the mean (s.e.m.) calculated from n = 3 independent experiments. Complete kinetics including all tested intronistat B concentrations are reported in **Figure S1**.

Besides informing on the overall potency of the compound, our assay allowed us to discriminate the effects of the compound on the first *vs* the second steps of splicing, because the *O. iheyensis* group IIC intron produces linear intron-3’-exon intermediates (I-3E) and free introns (I) that can be resolved by electrophoresis. Our data proved that in the presence of intermediate intronistat B concentrations (10-50 μM), the first step of splicing is ∼5-fold slower than in the absence of the compound (k_1_ = 0.037 ± 0.002 min^-1^ without compound, k_1_ = 0.007 ± 0.002 min^-1^ at 50 μM intronistat B, **Table S1**). Unexpectedly, we additionally noticed that intronistat B induces accumulation of I-3E, suggesting that the intron is unable to progress on to the second step of splicing. Indeed, our kinetic analysis proved that the second step of splicing is 34-fold slower in the presence *vs* absence of the compound (k_2_ = 0.031 ± 0.003 min^-1^ without compound, k_2_ = 0.001 ± 0.000 min^-1^ at 50 μM intronistat B, **Table S1**). These data indicate that intronistat B selectively inhibits the second step of splicing 7-fold more potently than the first step (**Figures 1 and S1**). Importantly, the splicing defects induced by intronistat B are comparable to those caused by active site mutations designed to impair the transition from the first to the second step of splicing, i.e. the so-called G- and U-triple mutants that inhibit active site protonation or the C377G mutant which impair active site toggling (Manigrasso et al., 2020; Marcia and Pyle, 2012).

In summary, our enzymatic studies establish that intronistat B displays a broader spectrum than previously suggested. It inhibits bacterial group IIC introns, which splice through a hydrolytic mechanism and produce linear introns, with a similar IC_50_ and a 5-fold higher *K*_i_ than fungal group IIB introns, which splice through a transesterification mechanism producing lariat introns. Moreover, our results show that intronistat B inhibits both the first and the second steps of group II intron splicing with a stronger effect on the second step. This unforeseen discovery could mean that intronistat B binds at or near the splice site with unexpected selectivity.

### Principles of molecular recognition between the conserved group II intron active site and splicing modulators

To visualize the exact binding mode and characterize the inhibitory mechanism of intronistat B at high resolution, we first solved the co-crystal structure of intronistat B bound to a previously-described construct of the *O. iheyensis* I1 intron (Marcia and Pyle, 2012), which encompasses its structural domains 1–5 (OiD1-5), in the exon-free state and in the presence of Mg^2+^ and K^+^ ions.

Our structure, determined at a resolution of 3.0 Å, shows that in the presence of the compound, the intron maintains an overall folded structure, similar to that of the compound-free intron (PDB entry 4E8M), with a root-mean-square deviation (RMSD) = 0.5 Å between the two states (**Table S2**). In the intronistat B-bound state the intron active site is intact and adopts the catalytically required triple helix conformation anchored to the M1/M2/K1 metal ion cluster (Marcia and Pyle, 2012).

To localize the compound in the crystallographic electron density, we calculated a simulated-annealing electron density omit-map. The map shows an extended positive peak in the active site (highest contour value = 4.5 σ, volume of the peak at 3.0 σ = 449 Å^3^), compatible with the chemical structure and molecular volume of intronistat B (predicted volume of intronistat B = 333 Å^3^, **Figure 2A-B**). Here, intronistat B makes specific contacts through its pyrogallol group with both M1 (2.1 Å from intronistat B O24 and O26 atoms) and M2 (2.1 Å from intronistat B O22 and O24 atoms, **Figure S3A**). As shown by our quantum-level calculations, this binding mode is possible, because intronistat B is deprotonated at the hydroxyl group in *para* in the pyrogallol moiety (pK_A_ = 5.7), analogously to other similar chelating groups (Credille et al., 2018). The pyrogallol moiety makes further hydrogen bond interactions with evolutionarily-conserved active site nucleotides, i.e. U375 (2.9 Å between intronistat B O26 atom and U375 OP1 atom), C358 (2.2 Å between intronistat B O22 atom and C358 OP1 atom) and G359 (3.1 Å between intronistat B O22 atom and G359 OP2 atom, **Figure S3A**). Additionally, intronistat B establishes a sequence-specific hydrogen bond with exon binding site 1 (EBS1), the motif that recognizes the exon sequence and thus defines the splicing site (3.2 Å between O18 in intronistat B and N1 in A181, **Figures 2A and S3A**). In this conformation, the real-space correlation coefficient (RSCC) calculated for intronistat B after refinement is 0.94, indicating very close similarity between the experimental and theoretical electron-density map for the compound (Pearce et al., 2017).

**Figure 2.**
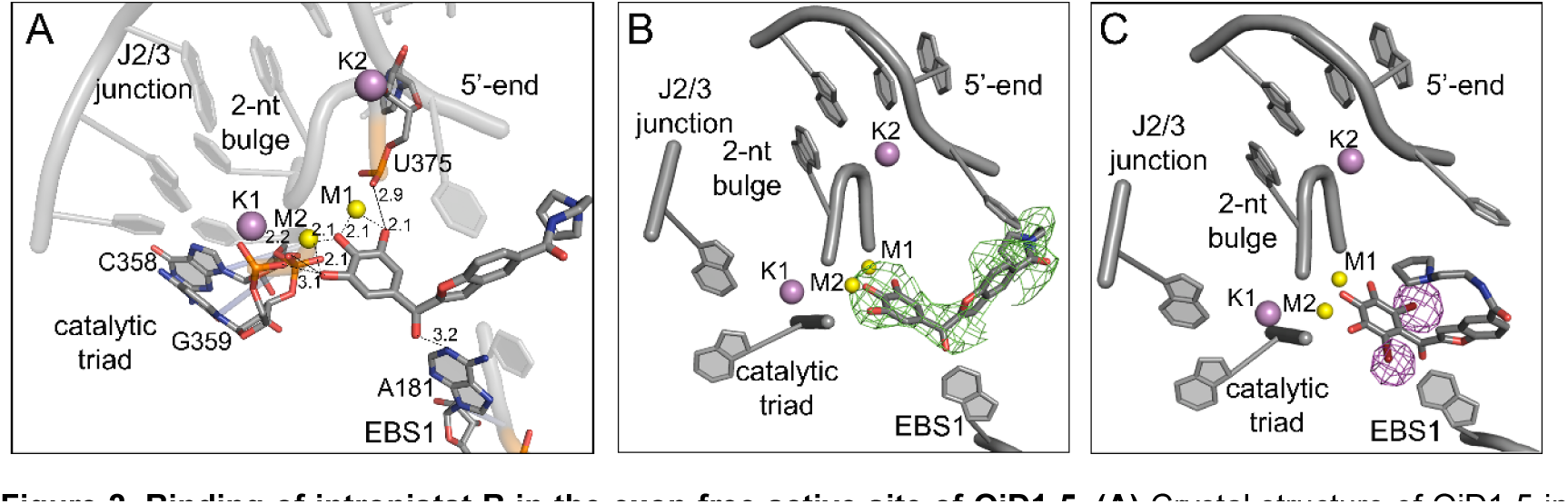
Binding of intronistat B in the exon-free active site of OiD1-5. **(A)** Crystal structure of OiD1-5 in the presence of Mg_2+_ (yellow spheres), K_+_ (purple spheres) and intronistat B (grey sticks). The interactions of intronistat B with active site elements are shown as black dotted lines (distances in Å). **(B)** Same active site as in panel A. Here, the F_o_-F_c_ simulated-annealing omit map obtained omitting intronistat B and contoured at 3σ is depicted in green mesh. **(C)** Crystal structure of OiD1-5 in the presence of Mg_2+_ (yellow spheres), K_+_ (purple spheres) and the di-brominated intronistat B derivative ARN25850 (grey sticks, RSCC = 0.88). The violet mesh represents the anomalous difference Fourier map contoured at 5σ, revealing the position of the bromide atoms.

To further validate this highly-specific compound binding mode, we additionally synthesized a di-brominated intronistat B derivative (ARN25850, **Figure S4**). We proved that this compound inhibits both *S. cerevisiae* group IIB (IC_50_ = 6.0 ± 0.9 μM) and *O. iheyensis* group IIC introns (K_i_ = 48.39 ± 17.09 μM, **Figure S4 and Table S1**). We then solved the crystal structure of OiD1-5 in complex with di-brominated intronistat B at a resolution of 2.8 Å (RMSD = 0.6 with respect to the crystal structure of intronistat B-bound OiD1-5). The crystallographic data, collected at λ = 0.92 Å, a wavelength where bromide displays an anomalous scattering coefficient *f”* = 4 electrons, enabled us to precisely localize the bromide atoms (Tiefenbrunn, 2014). The anomalous difference Fourier electron density map derived from our crystallographic data set shows two pronounced peaks within the intron core (highest contour value = 9.2 σ and 11 σ, respectively), at the expected *ortho* positions on the pyrogallol moiety (**Figure 2C**). This structure thus confirms unequivocally our modeling of the intronistat B conformation at the evolutionarily-conserved group II intron active site.

We further confirmed the specificity of the interaction by two independent ∼500 ns-long molecular dynamics (MD) simulations of the intronistat B-bound OiD1-5 structure, which show that the inhibitor stably maintains the binding pose captured crystallographically (RMSD_RNA_ = 5.23 ± 0.36 Å; RMSD_(intronistat B)_ = 1.54 ± 0.12 Å, **Figure S5**). In this position, the ligand mimics the splicing substrate (i.e. the scissile nucleotide) and forms a pseudo-Michaelis-Menten complex at the intron active site, stabilizing the M1-M2 internuclear distance at a value compatible with catalysis (d_M1-M2_ = 4.01 ± 0.46 Å; for reference in PDB entry 4E8M d_M1-M2_ = 3.7 Å, **Figures 3 and S5-6**).

**Figure 3.**
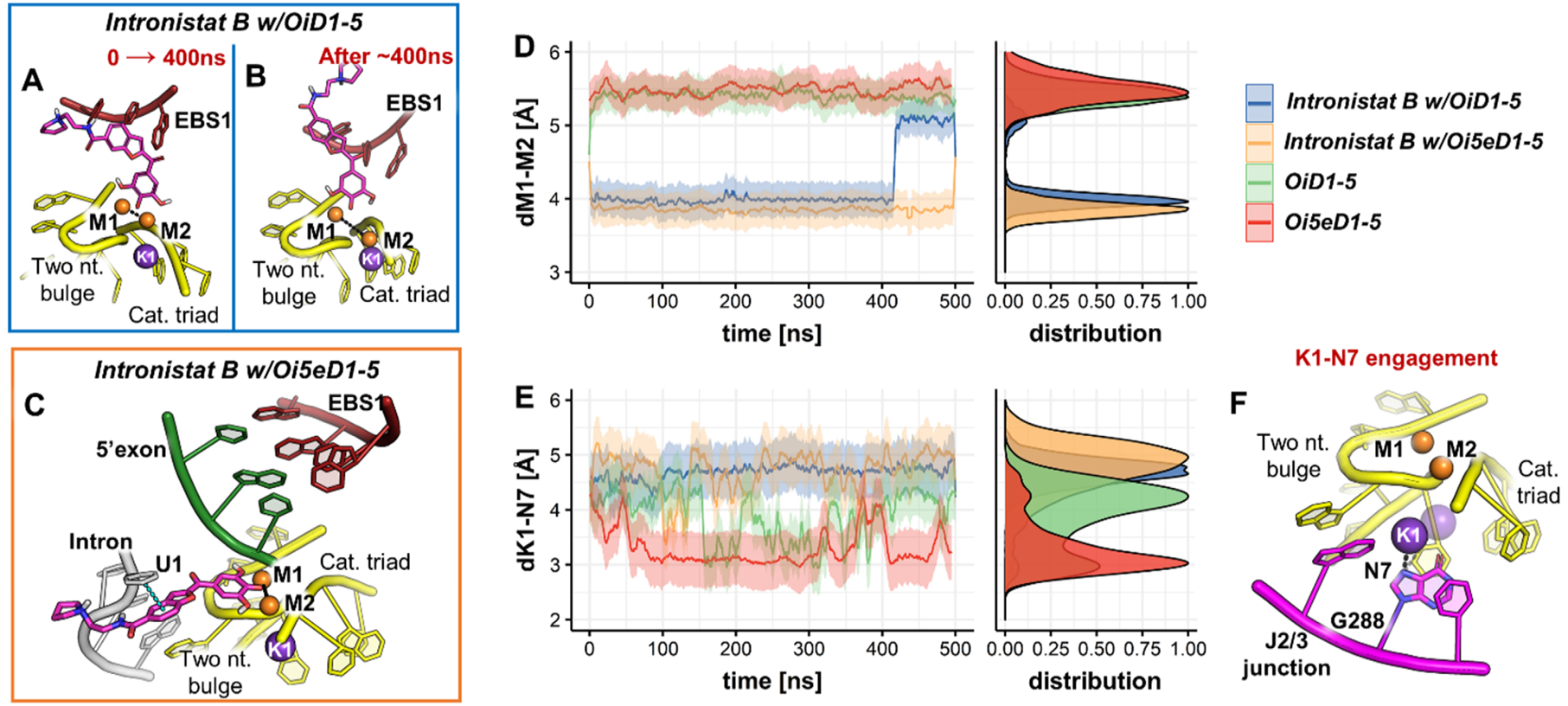
MD simulations. **(A)** Representative binding mode of intronistat B as found during the first 400 ns of MD simulations – when bound to the free intron. The compound (purple sticks) coordinates the two catalytic magnesium ions M1-M2 (orange spheres), thus competing with the splice junction and mimicking a pseudo Michaelis-Menten complex. The catalytic triad (yellow), the two-nucleotide bulge (yellow) and the exon binding site 1 (EBS1, red) residues are shown as cartoon. **(B)** Representation of the alternative binding mode of intronistat B captured by MD simulations when bound to the free intron. Here, the compound loses the interaction with M2 while preserving the coordination with M1. The representation follows that of panel A. **(C)** MD simulations show that, in the presence of the 5’exon, intronistat B coordinates both the catalytic M1-M2 ions and engages one additional π-π interaction with the U1 nucleotide of the intron. The representation follows that of panel A. **(D)** The internuclear distance between the catalytic M1-M2 ions is reported as a function of simulation time. As expected, in both the free (green) and 5’exon-bound intron (red) apo-system the metals are stabilized at a distance not compatible with catalysis. In the presence of intronistat B, in both the free (blue) and the 5’-exon-bound (yellow) intron systems the two metals are coordinated by the compound (panel A and C, respectively), mimicking a pseudo-Michaelis Menten complex even in the absence of the native substrates, thus rigidifying the active site in a non-productive conformation. **(E)** The distance between K1 and the atom N7 of G288 is plotted as a function of simulation time. In the absence of intronistat B, the K1-N7 interaction is engaged in both the free (green) and 5’exon-bound (red) intron systems. When intronistat B coordinates the catalytic M1-M2 ions (panel D), it alters the functional dynamics of the active site preventing the formation of K1-N7 interaction in both the free and 5’-exon-bound system, ultimately impairing the splicing initiation and progression. **(F)** The catalytic K1 ion (purple sphere) engages the interaction with N7 atom of G288 (purple sticks) at the J2/3 junction (purple cartoon). If the ion is not in place (purple semitransparent sphere), the K1-N7 interaction is not established, ultimately inhibiting the catalytic activation of the intron for the first splicing step, and its structural rearrangement necessary for splicing progression (Manigrasso et al., 2020). The representation follows that of panel A.

Taken together, our OiD1-5 structures with intronistat B and its brominated derivative, and the corresponding MD simulations, demonstrate that intronistat B inhibits splicing because it binds to the active site of group II introns, where it forms direct, specific interactions with conserved catalytic nucleotides and metal ions.

### The mechanism of inhibition of the first step of splicing

Established that intronistat B binds at the active site of group II introns and inhibits different catalytic steps with different potency, we sought to better understand the underlying mechanism of such unexpected selectivity. To this end, we solved co-crystal structures of *O. iheyensis* group IIC intron in complex with intronistat B at each step of the splicing cycle. For this study, besides the OiD1-5 construct described above, we additionally used the previously-described Oi5eD1-5 intron, which comprises the short 5’-exon sequence UAUU at its 5’-terminal end (Marcia and Pyle, 2012).

Using Oi5eD1-5, we first obtained a structure of the pre-catalytic state (the stage prior to the first step of splicing) in the presence of Ca^2+^, K^+^, and intronistat B at 4.0 Å resolution (**Table S2**). The simulated-annealing electron density map generated by omitting the G1 at the 5’ splice junction demonstrates that in this structure the active site is occupied only by the reaction substrates, i.e. the 5’-splice junction and the reaction nucleophile. Despite the presence of high concentrations of intronistat B in the crystallization buffer (1 mM, 588-fold over the K_i_), this structure adopts the exact same conformation reported for the corresponding apo-form (RMSD = 0.6 Å with respect to PDB id: 4FAQ, **Figure S7A**).

We also determined a co-crystal structure at 3.1 Å resolution of Oi5eD1-5 and intronistat B in the state immediately following the first step of splicing, which is obtained in the presence of Mg^2+^ and K^+^. Analogously to the structure obtained in the presence of calcium, also the structure obtained in the presence of magnesium shows the compound-free intron, in which the scissile phosphate and the 5’-exon coordinate M1 and M2 in the same conformation reported for the corresponding apo-form (RMSD = 0.4 Å with respect to PDB id: 4FAR, **Figure S7B**). These two structures of Oi5eD1-5 suggest that under the experimental crystallization conditions (100 mM Mg^2+^, 30°C) the splice junction outcompetes intronistat B from the active site. We had previously established, however, that immediately after 5’-exon hydrolysis these crystallization conditions trap the scissile phosphate in an unproductive conformation, which is unlikely to be of physiological relevance (Manigrasso et al., 2020).

To mimic a more physiological conformation of the active site immediately after 5’-exon hydrolysis, we thus co-crystallized OiD1-5 (the free intron) with the 5’-exon-like oligonucleotide 5’-AUUUAU-3’, and we solved the X-ray structure of this complex in the presence of intronistat B and of Mg^2+^ and K^+^ at 3.3 Å resolution. This structure reveals a well-folded intron possessing an intact catalytic site that adopts the expected triple helix conformation (RMSD = 1.0 Å with respect to the corresponding apo-form, PDB id: 4FAW, **Figure 4**). An omit map, generated before modelling the compound, displays a peak in the active site, compatible with the binding of intronistat B (highest contour value = 4.0 σ, volume of the peak at 3.0 σ = 301 Å^3^). Unexpectedly, this volume is oriented differently from the volume occupied by intronistat B in the exon-free state (RSCC = 0.90, **Figure 2B**). The group II intron active site still recognizes the compound’s pyrogallol motif through M1 (1.9 Å from intronistat B O22 atom and 2.1 Å from intronistat B O24 atom), M2 (1.9 Å from intronistat B O24 atom and 2.2 Å from intronistat B O26 atom), and its conserved catalytic residues, i.e. C377 (3.2 Å between intronistat B O22 atom and C377 OP1 atom, 2.6 and 2.3 Å between intronistat B O24 atom and C377 OP1 and OP2 atoms and 3.0 Å between intronistat B O26 atom and C377 OP2 atom), U375 (2.6 Å between intronistat B O22 atom and U375 OP1 atom), G359 (2.8 Å between intronistat B O24 atom and G359 OP2 atom), and C358 (2.4 Å between intronistat B O26 atom and C358 OP2 atom). Interestingly, in this structure, the nucleotides around the splice junction also contribute to the recognition of the compound, i.e. U0 [3.3 Å between intronistat B O22 atom and U0 O3’ atom and 3.1 Å between intronistat B O24 atom and U0 O3’ atom] and U2 (weak 3.9 Å contact between O18 of intronistat B and O4’ of U2). However, in this structure intronistat B has lost its interaction with A181 in the EBS1 site (15 Å between O18 in intronistat B and N1 in A181), which is now based-paired to its cognate 5’-exon nucleotide U0 (**Figures 4 and S3B**).

**Figure 4.**
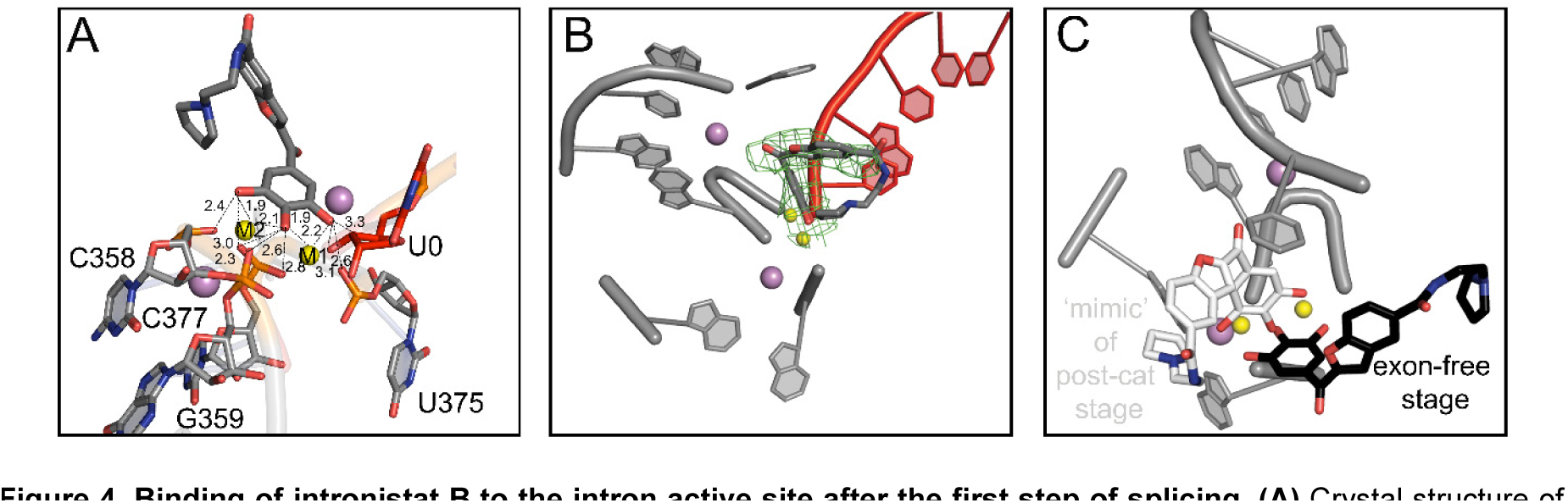
Binding of intronistat B to the intron active site after the first step of splicing. **(A)** Crystal structure of OiD1-5 in the presence of Mg^2+^ (yellow spheres), K^+^ (purple spheres), the 5’-exon-like oligonucleotide 5’-AUUUAU-3’ (red sticks), and intronistat B (grey sticks). Interactions of intronistat B with active site elements are shown as black dotted lines (distances in Å). **(B)** F_o_−F_c_ electron density omit map for the structure reported in panel A, calculated by omitting intronistat B, contoured at 3σ, and represented as a green mesh. **(C)** Superposition of the intronistat B binding mode in the cleaved state (white sticks, from panel A) and in the exon-free state (black sticks, from **Figure 2**).

**Figure 5.**
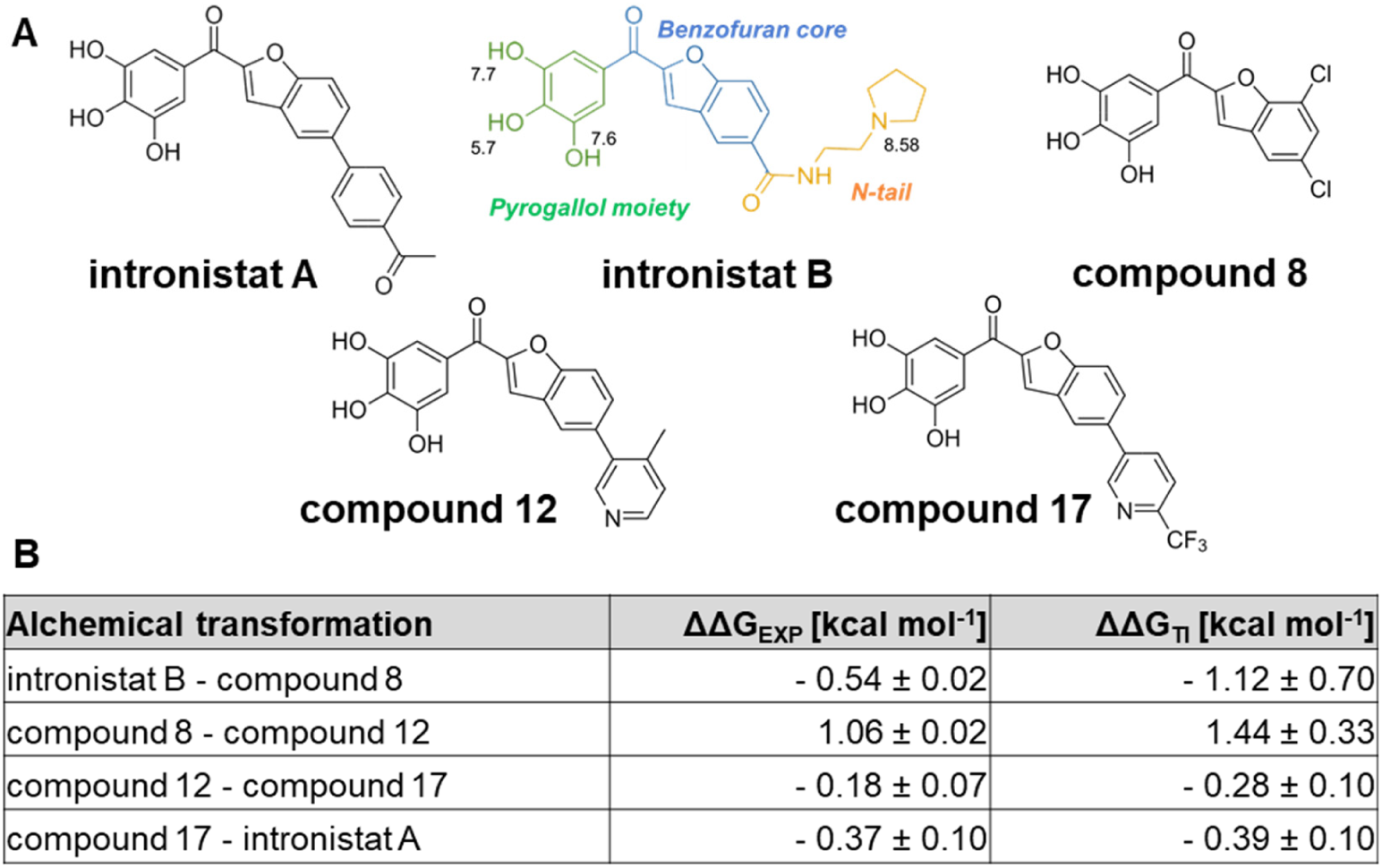
Splicing modulators. **(A)** Chemical structures of the splicing modulators described in the text. The chemical groups of intronistat B are highlighted in green (pyrogallol moiety), blue (benzofuran core), and yellow (N-tail). The atom number and the corresponding pK_a_ values are also reported. **(B)** Relative binding free energy of splicing modulators. The ΔΔG values computed from experimental *K_i_* (ΔΔG_EXP_) as well as that estimated computationally (ΔΔG_TI_) are reported for each alchemical transformation.

Two additional independent ∼500 ns-long MD simulations of intronistat B bound to OiD1-5 and 5’-exon show that the compound is steadily anchored at the active site as shown by X-ray crystallography (RMSD_RNA_ = 3.81 ± 0.22 Å; RMSD_(intronistat B)_ = 1.82 ± 0.17 Å, **Figure S6**). Similar to the free intron structure, also in the presence of the 5’-exon the two catalytic M1-M2 ions coordinate the compound and are stabilized at a distance d_M1-M2_ = 4.01 ± 0.46 Å (**Figure 3D**). Additionally, during our simulations, the contact between U2 and intronistat B is reinforced by the formation of one additional π-π stacking interaction between the U2 nucleobase and the benzofuran core of the ligand (dπ_U2_-π_intronistat B_ = 3.91 ± 1.22 Å, **Figure 3C**), which further stabilizes the binding of the compound in its crystallographic pose.

Altogether, these new structures and extensive MD simulations of the group II intron in the presence of intronistat B before and after the first step of splicing suggest that the compound competes with the scissile phosphate to inhibit this catalytic step, curiously adopting a different binding conformation than in the exon-free state. We therefore decided to probe this distinctive conformation, coupled to the inhibition constants of intronistat B and previously-reported analogues, by computing the relative binding free energy (ΔΔG) of these compounds at the intron active site [**Figure S8**, (Cournia et al., 2017; Klimovich et al., 2015)]. To this aim, we first used the compound-and 5’-exon-bound structure of OiD1-5 (**Figure 4**). Considering the structural homology between our *O. iheyensis* and the ai5γ introns, we then correlated these relative binding free energy values to the respective experimental inhibition constants of the compounds. We first compared intronistat B (*K*_i_ = 0.36 ± 0.02 μM) with compound 8 [APY083, *K*_i_ = 0.90 ± 0.10 μM (Fedorova et al., 2018)]. Our alchemical free energy calculation estimated a difference in binding free energy between compound 8 and intronistat B ΔΔG_(compound 8 – intronistat B)_ = -1.12 ± 0.70 kcalꞏmol^-1^ (**Figures 5 and S9-10**). Importantly, this ΔΔG value is in line with the ΔΔG that can be derived from the experimental *K*_i_ of the two compounds *via* the Gibbs equation. Thus, the crystal structure provides an accurate model for intronistat B binding, indicating that intronistat B analogues bind homologous group II introns in the same manner (ΔG_introninstat B_ = -8.85 ± 0.01 kcalꞏmol^-1^, ΔG_compound 8_ = -8.31 ± 0.20 kcalꞏmol^-1^, ΔΔG_(compound 8 – intronistat B)_ = -0.54 ± 0.10 kcalꞏmol^-1^).

To further validate this distinctive binding pose, we then compared three additional pairs of previously-reported intronistat B analogs [compound 8 *vs* compound 12 (APY081), *K*_i(12)_ = 5.3 ± 0.2 μM, ΔΔG_(8-12)_ = 1.063 kcalꞏmol^-1^; compound 12 (APY081) *vs* compound 17 (APY097), *K*_i(17)_ = 3.9 ± 0.5 μM, ΔΔG_(12-17)_ = -0.18 ± 0.54 kcalꞏmol^-1^; compound 17 (APY097) *vs* intronistat A (APY101), *K*_i(intronistat A)_ = 2.1 ± 0.2 μM, ΔΔG_(17-intronistat A)_ = -0.37 ± 0.54 kcalꞏmol^-1^, **Figures 5 and S11-12** (Fedorova et al., 2018)]. Remarkably, also these ΔΔG estimates overlap well with the ΔΔG derived from the experimental *K*_i_ (ΔΔG_(8-12)_ = 1.44 ± 0.33 kcalꞏmol^-1^; ΔΔG_(12-17)_ = -0.28 ± 0.1 kcalꞏmol^-1^; ΔΔG_(17-intronistat A)_ = -0.39 ± 0.1 kcalꞏmol^-1^).

The close correspondence between the experimental and computationally estimated ΔΔG values suggest that the intronistat B binding mode captured by our crystal structures accounts for all major interactions responsible for the experimentally-measured *K*_i_ of the ligands.

### The mechanism of inhibition of the second step of splicing

To gain further insights into the preferential inhibition of the second step of splicing by group II intron inhibitors, we solved the crystal structure of OiD1-5 at 3.6 Å resolution, in the presence of intronistat B and of Mg^2+^ and Na^+^ ions. This condition forces the intron active site into a transiently inactive, so-called “toggled” conformation, which is achieved *via* rotation of residues 287-289 of J2/3 junction around their backbone, and which is mechanistically important because it favors the transition from the first to the second step of splicing (Manigrasso et al., 2020; Marcia and Pyle, 2012).

In our new sodium structure with intronistat B, the intron retains the same overall structure as in the apo form (RMSD = 1.2 Å with respect to PDB id: 4FAX). In this structure, the F_o_-F_c_ electron density map, generated before modelling the compound, displays a peak in the active site, which is absent in the apo form, and which is compatible with the size of the pyrogallol moiety of intronistat B (highest contour value = 4.3 σ, volume of the peak at 3.0 σ = 46 Å^3^, RSCC = 0.92, **Figure 6**). This density suggests that intronistat B is bound to the active site also in the presence of sodium, albeit in a more flexible conformation than for the structures with potassium (see **Figures 2 and 4**). Most importantly, a simulated-annealing omit map obtained by omitting residues 287-289 of J2/3 junction and M1/M2, besides intronistat B, unequivocally reveals that the intron adopts a triple helix and not the expected toggled conformation (**Figure 6C**). In line with this observation, the catalytic ions M1 and M2, which are normally released from the active site in the toggled state, are instead still present in our structure. M1 is located at 2.0 Å from intronistat B O26 and at 2.1 Å from intronistat B O24, while M2 is at 2.0 Å from intronistat B O24 and 2.1 Å from intronistat B O22. Intronistat B makes further hydrogen bond interactions with G374 (3.3 Å between intronistat B O26 and G374 O3’), C358 (2.8 Å between intronistat B O22 and C358 OP1), G359 (2.3 Å between intronistat B O24 and G359 OP1), C377 (2.3 Å between intronistat B O24 and C377 OP2), and A181 (3.2 Å between O18 in intronistat B and N1 in A181) (**Figure 6A**).

**Figure 6.**
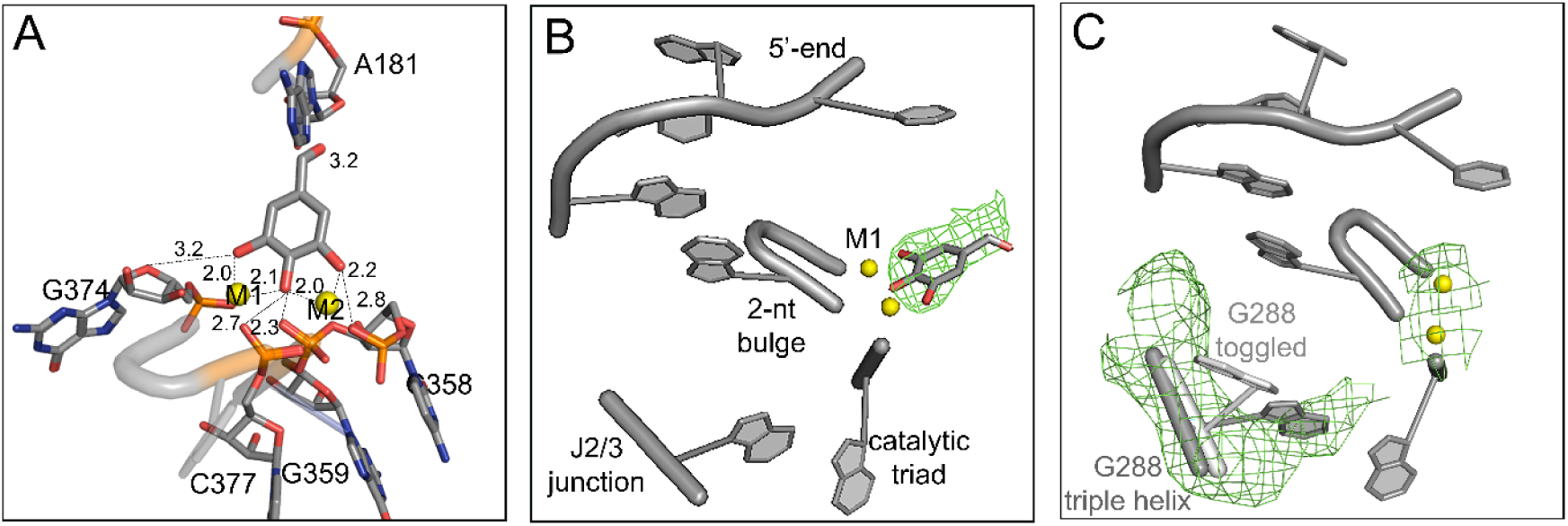
Intronistat B prevents the toggling of the active site. **(A)** Crystal structure of OiD1-5 in the presence of Mg_2+_ (yellow spheres), Na_+_, and intronistat B (grey sticks, only the pyrogallol moiety is visible in the electron density in this structure, see panel B). Interactions of intronistat B with active site elements are shown as black dotted lines (distances in Å). **(B)** F_o_−F_c_ electron density map for the structure reported in panel A generated before modelling intronistat B and shown as a green mesh and contoured at 2.9σ. **(C)** F_o_−F_c_ simulated annealing electron density omit map (green mesh) for the structure reported in panel A, calculated by omitting intronistat B, the J2/3 junction residues and both M1 and M2 and contoured at 2.7σ. This omit map shows that the J2/3 junction adopts the triple-helix conformation and not the toggled conformation. For reference, the toggled conformation is modeled here as light grey sticks from PDB id: 4FAX.

While the flexibility of intronistat B prevents us from simulating the dynamics of the sodium structure, our equilibrium MD simulations of the OiD1-5 and Oi5eD1-5 structures with and without intronistat B in the presence of potassium provide a precise mechanistic explanation for how the compound inhibits active site toggling. Specifically, multiple ∼500 ns-long MD replicas show that the presence of intronistat B prevents the formation of the catalytically-essential interaction between the active site potassium ion K1 and the N7 atom of the conserved G288 nucleotide [**Figure 3F**, (Manigrasso et al., 2020)]. In the absence of intronistat B the K1-N7 interaction is properly established (d_K1-N7_ = 3.12 ± 0.43 Å and d_K1-N7_ = 3.07 ± 0.52 Å for OiD1-5 and Oi5eD1-5, respectively, **Figure 3E**). Instead, in the presence of intronistat B the active site dynamics are altered, preventing the engagement of K1 with G288 N7 (d_K1-N7_ = 4.72 ± 0.53 Å and d_K1-N7_ = 4.68 ± 0.61 Å, for OiD1-5 and Oi5eD1-5, respectively, **Figures 3E-F and S5-6**).

In summary, by impeding the establishment of the K1-N7 interaction and by preventing active site toggling, as shown by our Na^+^ structure and MD simulations, intronistat B de-activates the intron and impairs its transition from the first to the second step.

## DISCUSSION

Splicing is a vital and ubiquitous reaction that ensures the correct maturation of transcribed genes in all forms of life. Its modulation by small molecule compounds has recently emerged as a very promising therapeutic strategy to treat pathogenic infections, as well as human genetic diseases and cancer, but the principles by which splicing modulation is achieved are still largely unclear at the molecular level (Fedorova et al., 2018; Lee and Abdel-Wahab, 2016; Naryshkin et al., 2014). In this context, our 6 new crystal structures, corroborated by enzymatic data and microsecond-long atomistic simulations, now show at atomic resolution how the evolutionarily-conserved active site of a splicing ribozyme interacts specifically and selectively with small molecule modulators (**Figure 7**). Nearly 40 years after establishing that the active site of the ribosome recognizes specific small molecule inhibitors (Brodersen et al., 2000; Moazed and Noller, 1987; Wilson, 2009), our results exemplify the power of using RNA-binding molecules to mechanistically probe and functionally modulate another fundamental biological reaction catalyzed by RNA, splicing.

**Figure 7.**
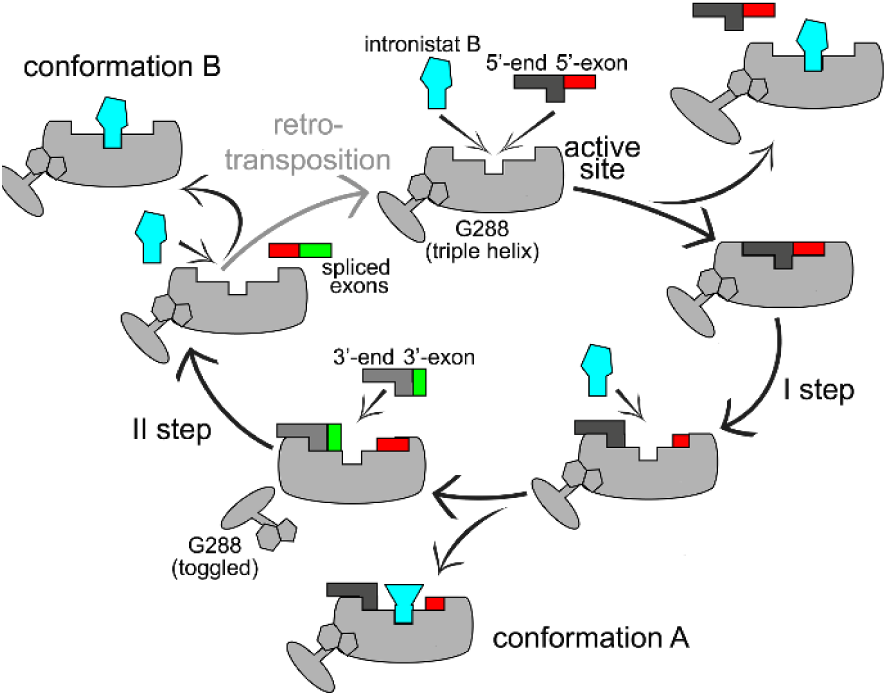
Schematic mechanism of group II intron splicing inhibition by small molecule compounds. In the pre-catalytic state, intronistat B competes with the 5’-splice junction at the intron active site to inhibit the first step of splicing. Intronistat B also binds the intron active site in the presence of the 5’-exon after the first step of splicing, by establishing only sequence-unspecific interactions and contacts with active site metal ions (conformation A, **Figure 4**). In this state, intronistat B prevents the intron from switching from its triple-helix to its toggled conformation, thus inhibiting the second step of splicing more potently than the first. Finally, intronistat B also binds the exon-free intron by establishing contacts with the catalytic metal ions and both sequence-specific and sequence-unspecific interactions (conformation B, **Figure 2**).

The highly-structured splicing site is perfectly suited to tightly capture and specifically recognize small organic compounds through a complex interaction network established by conserved nucleotides and the catalytic metal ion cluster. Unexpectedly, at the splicing site, the compounds display two distinct mechanisms of action. First, they compete with the substrates of the splicing reaction, i.e., the splice junctions. Second, they stabilize the intron active site in its so-called triple helix configuration, mimicking the formation of an unproductive pseudo Michaelis-Menten complex in the absence of the native substrates. This binding mode prevents the formation of the catalytically essential K1-N7 interaction at the active site, ultimately impairing the intron from toggling into an open and transiently-inactive state after completion of the first step of splicing. Through mutagenesis, metal ion replacement approaches, and atomistic simulations, we had previously demonstrated that such transiently-inactive toggled configuration is essential for preparing the intron active site for the second step of splicing (Manigrasso et al., 2020; Marcia and Pyle, 2012). In line with such mechanistic data, here we observe that by impeding active site toggling, intronistat B selectively induces a severe defect of the second step.

As revealed explicitly for intronistat B, the compound’s interactions with the intron are primarily sequence-unspecific, in line with the broad spectrum of the compound, which inhibits both bacterial and organellar introns belonging to different classes (groups IIB and IIC) and following different splicing pathways (hydrolysis *vs* transesterification). Nevertheless, in the exon-free state, intronistat B also establishes sequence-specific contacts with a functional nucleotide (A181 in the EBS1 site). This residue is not conserved in group II introns and it can be replaced by purines or by pyrimidines, i.e. a G in *Escherichia coli* EcI5 group IIB intron (Monachello et al., 2021) and in *Lactococcus lactis* LlI1 group IIA intron (Belhocine K, 2007), a C in *Thermosynechococcus elongatus* Tel4h group IIB intron (Haack DB, 2019) and *Bacillus halodurans* BhI1 group IIC intron (Toor N, 2006), or a U in *Pylaiella littoralis* PliLSUI2 group IIB intron (Wiryaman and Toor, 2017). This observation suggests that a small molecule like intronistat B can be designed and modified to establish stronger, sequence-specific contacts with the EBS1 site to generate more selective, species-specific compounds. In this context, our computational protocol to estimate the relative binding free energy of intronistat B analogues can now be used for the rational design of such sequence-specific modulators, building upon the intronistat B-intron interaction network captured crystallographically in our work.

Quite remarkably, intronistat B not only binds tightly to the active site and displays a selective dual mechanism of inhibition, but it also dynamically adapts to conformational changes in the intron active site, by changing its binding mode while the intron progresses through its different catalytic states. At a time when the determinants that govern RNA-small molecule interactions are still largely uncharacterized, our observations of the exquisite selectivity and specificity of intronistat B showcase the potential of modulating RNA structure-functional dynamics very precisely with small molecules.

Our data thus represent a solid experimental basis for directing the rational design of splicing modulators through structure-based strategies, in the same way that is traditionally applied to protein design and drug discovery. Mechanistic studies on the mode of action of ribosomal RNA inhibitors provided important insights for developing new species-specific antimicrobial agents overcoming the insurgence of resistance, besides crucially improving our understanding of the process of translation (Wilson, 2009). Similarly, our study could help improve the potency of intronistat B, which is still insufficient to make this tool compound a suitable drug lead for the treatment of fungal infections (low micromolar K_i_ and IC_50_ values). Potentially, group II intron inhibitors could also have relevance in the context of bacterial infections. In bacteria, group II introns are often located in plasmids associated to genes that confer antibiotic resistance (Ichiyanagi K, 2003; Partridge SR, 2018). Although a clear connection between group II intron activity and bacterial infectivity is still lacking, it cannot be excluded that inhibiting bacterial splicing could prevent antibiotic resistance. Certainly, though, bacterial group II introns act as retroelements (Dai and Zimmerly, 2002) and thus have established biotechnological applications in genome editing, as so-called “targetrons” (Enyeart PJ, 2014). As such, bacterial group II introns can be used to site-specifically deliver cargo genes into genomic sites, including genes for fluorescent proteins, phage resistance, and antigens. However, targetrons have limited applicability because of their off-target effects and poor specificity. By establishing that the exon-free state of group II introns (i.e. the state that catalyzes retro-homing) binds small molecules tightly and in a sequence-specific manner, our study now opens up the possibility of developing improved targetrons. For instance, it could be envisaged to design targetrons that are constitutively inhibited by photoactivatable intronistat B derivatives, deliver these ribozymes systemically, and only photoactivate them in the desired target tissues, thus reducing their toxicity (Bayley et al., 1987; Ellis-Davies, 2007). Most importantly, by proving that the conserved RNA-core of self-splicing ribozymes is capable of recognizing small molecules specifically and selectively, our work sets an unprecedented and unexpected solid rationale for the design of analogous compounds to also inhibit the eukaryotic spliceosome. The spliceosome active site derives evolutionarily from that of group II introns and preserves its key structural, chemical and functional properties (Marcia et al., 2021). Not surprisingly, by superimposing the intron and spliceosome active site structures, we note that compounds analogous to intronistat B could fit within the catalytic core of spliceosomal complexes formed at different stages of catalysis, including the B* complex (i.e. before the first step of splicing, PDB id: 5Z56, 5Z57, 5Z58) and the C_i_ complex (i.e. after the first step of splicing, PDB id: 7B9V). Here, the compounds could anchor at the M1-M2-K1 site with their pyrogallol or an analogous moiety, as our work now proves possible. In this binding pose, the compounds would be able to explore multiple conformations, interacting with catalytic nucleotides, metal ions or protein splicing factors (**Figure S13**). Importantly, by binding at the active site, the compounds would also necessarily reach close to nucleotides around the splice junctions. Establishing direct interactions with pre-mRNA nucleotides close to the splice junctions would crucially offer unprecedented opportunities for sequence-specific and gene-selective modulation of splicing. Such selectivity cannot be achieved by targeting splicing assembly factors, a splicing modulation strategy that is currently widely explored, but that inevitably causes severe toxicity (Lee and Abdel-Wahab, 2016; Seiler et al., 2018). Importantly, achieving gene-selective splicing modulation with small molecule inhibitors could also offer advantages over the established use of splice-switching oligonucleotides (SSO), which are expensive, display poor cellular uptake and tissue/cell-specificity, and are prone to off-target effects (Taylor et al., 1999). More broadly, considering that the group II intron and spliceosome active site architecture and catalytic mechanism are also very similar to those of nucleic acid-processing protein enzymes, like DNA/RNA polymerases and endo/exonucleases, it should not be excluded that the design of specific inhibitors against one of these convergently-evolved enzymes could then inform the rational design of analogues that inhibit other target classes (Genna et al., 2018; Genna et al., 2019; Manigrasso et al., 2021a).

In summary, our data demonstrate that the conserved RNA-based active site of splicing enzymes can specifically and selectively recognize small molecule modulators. These RNA-targeting compounds thus emerge as very informative tools to mechanistically probe a vital biological reaction, such as splicing, and to foster the design of new biotechnological, genetic engineering and pharmacological applications.

## ACKNOWLEDGEMENTS

We thank the scientists at the ID30-A1 and ID30B ESRF beamlines for support with data collection, Dr Isabel Chillon (CNRS, IGMM Montpellier) for help with initial splicing kinetic assays, and Ombeline Pessey and Ines Dieryck for excellent technical assistance. We thank Prof Annalisa Pastore, Prof Kristina Djinovic-Carugo, Dr Isabel Chillon, Michela Nigro and Shekhar Jadhav for critical reading of the manuscript. We also thank all members of the Marcia and De Vivo labs for helpful discussion. Work in the Marcia lab is partly funded by ITMO Cancer (18CN047-00), Région Auvergne Rhône Alpes (project R21105CC; allocation RPH21004CCA), FINOVI (AAP15), Canceropole CLARA (Oncostarter), and by the Fondation ARC pour la recherche sur le cancer (PJA-20191209284). The Marcia lab uses the platforms of the Grenoble Instruct Center (ISBG: UMS 3518 CNRS-CEA-UJF-EMBL) with support from FRISBI (ANR-10-INSB-05-02) and GRAL (ANR-10-LABX-49-01) within the Grenoble

Partnership for Structural Biology (PSB). MDV thanks the Italian Association for Cancer Research (AIRC) for financial support (IG 23679).

## DATA AVAILABILITY STATEMENT

Coordinates and structure factors have been deposited in the Protein Data Bank under accession codes 8OLS, 8OLV, 8OLW, 8OLY, 8OLZ and 8OM0.

## AUTHOR CONTRIBUTIONS

MM and MDV conceived and designed the work; IS determined all crystal structures and performed all enzymatic assays; JM performed all MD simulations; AA and NB synthesized ARN25850; all authors interpreted the data; IS, JM, MDV, and MM produced the first draft of the manuscript; all authors approved the final version of the manuscript. MM and MDV contributed equally.

## METHODS

### Cloning and mutagenesis

Four different constructs were used in this work. The wild-type *O. iheyensis* group II intron sequence (479 nt) flanked by its 5’ exon (223 nt) and by its 3’ exon (101 nt), corresponding to the previously-described pOiA and pOi5 constructs (Manigrasso et al., 2020; Marcia and Pyle, 2012; Toor et al., 2008), was used for the analysis of splicing kinetics. The previously-described OiD1-5 construct (Manigrasso et al., 2020; Marcia and Pyle, 2012; Toor et al., 2010; Toor et al., 2008) was used to crystallize the intron in the substrate-free state. The previously-described Oi5eD1-5 construct, corresponding to OiD1-5 following the 5’-exon 5’-UUAU sequence (Marcia and Pyle, 2012) was used to crystallize the intron in the pre-catalytic stage. A new crystallization construct corresponding to OiD1-5 without the three 5’-terminal Gs was used to crystallize the intron in the ’mimic’ of the post catalytic stage. The deletion was done by Gibson assembly using the Gibson Assembly Kit (NEB). The restriction enzyme ClaI (NEB) was used for linearization of pOiA templates, while BamHI (NEB) was used for linearization of all templates of the crystallization constructs. All constructs were confirmed by DNA sequencing (Eurofins).

### *In vitro* transcription and purification

All constructs were linearized by digestion with the appropriate endonucleases at 37°C overnight and transcribed *in vitro* using T7 polymerase (Marcia and Pyle, 2012). The constructs used for the crystallography experiments were then purified under native conditions (Chillon et al., 2015), buffered again and concentrated to 80 µM in 10 mM MgCl_2_ and 5 mM sodium cacodylate pH 6.5. The construct used for kinetic studies was radiolabeled during transcription, purified to a denatured state (Marcia and Pyle, 2012), and subsequently refolded (Toor et al., 2008).

### Splicing assays

Purified radiolabeled intron precursor was refolded by denaturation at 95 °C for 1 min in the presence of 40 mM Na-MOPS pH 7.5, and cooled at room temperature for 2 min. Then KCl to a final concentration of 150 mM and MgCl_2_ to a final concentration of 5 mM were added (Manigrasso et al., 2020; Marcia and Pyle, 2012), and the refolded intron was incubated with various concentrations of intronistat B [N-(2-(pyrrolidin-1-yl)ethyl)-2-(3,4,5-trihydroxybenzoyl)benzofuran-5-carboxamide; Sigma] or ARN25850 (self-synthesized, see below) at 37 °C. Aliquots at different time points were quenched with urea and analyzed on a 5% denaturing polyacrylamide gel (Daniels et al., 1996). The kinetic rate constants (k_obs_) in presence of each compound concentration were calculated using the Prism 8 package (GraphPad Software) and plotted against the inhibitor concentration according to the following equation to determine K_i_ values: *k*_obs_ = *k*_max_/(1+[I]/K_i_), where *k*_obs_ and *k*_max_ are the rate constants measured in the presence and in the absence of the inhibitor, respectively, [I] is the concentration of the inhibitor and K_i_ is the inhibition constant (Fedorova et al., 2018). Experiments were performed in triplicate. Data represent average ± s.e.m. The IC_50_ value was calculated by measuring the fraction of precursor (5e-I-3e) at the reaction time of 15 min (F_5e-I-3e, 15min_), calculating the percentage of reacted precursor at each concentration of compound according to the following equation: %_5e-I-3e_ = 100*(F_5e-I-3e, 15min, [I]max_ -F_5e-I-3e, 15min, [I]_)/(F_5e-I-3e, 15min, [I]max_ -F_5e-I-3e, 15min, DMSO_), and fitting the percentage of reacted precursor as function of the compound concentration according to the following function: %_5e-I-3e_ = 100/(1+[I]/IC_50_). Data are reported as average ± s.e.m.

### Crystallization

The natively purified intron was mixed with a 0.5 mM spermine solution and with the crystallization buffer in a 1:1:1 volume ratio (Marcia and Pyle, 2012). Crystals at the ‘mimic’ of the post-catalytic stage were crystalized in presence of a substrate oligonucleotide 5’-AUUUAU-3’ 100 μM. Crystals were grown at 30°C by the hanging drop vapor diffusion method using 2 μL sample drops and 300 μL crystallization solution in a sealed chamber (EasyXtal 15-Well Tool, Qiagen). Crystals were soaked for 1h (for the Na structure or with the dibromo Intronistat B derivative) or 2h 30’ (for the structures in the pre-catalytic, post-catalytic, ‘mimic’ of the post-catalytic and the exon-free stages) in a solution containing the corresponding crystallization buffers supplemented with 1 mM intronistat B or ARN25850 and cryo-protected with 25% ethylene glycol before flash frozen in liquid nitrogen. Crystals were harvested after 2 – 3 weeks, except those used to solve the structures in the prehydrolytic states, which were frozen within 24 h. The crystallization solutions used to solve the structures presented in this work were composed of: (1) 100mM K-Acetate, 100 mM KCl, 100 mM CaCl_2_, 50 mM Na-HEPES pH 7.0, 3% PEG 8000 for the calcium data set representing the pre-catalytic stage; (2) 100 mM Mg-Acetate, 150 mM KCl, 10 mM LiCl, 50 mM Na-HEPES pH 7.0, 4% PEG 8000 for the potassium data set representing the post-catalytic stage; (3) 100 mM Mg-Acetate, 200 mM KCl, 50 mM Na-HEPES pH 7.0, 6% PEG 8000 in the presence of the 6-mer RNA 5’-AUUUAU-3’ for the potassium data set representing the ‘mimic’ of the post-catalytic stage; (4) 100 mM Mg-Acetate, 150 mM NaCl, 50 mM Na-HEPES pH 7.0, 5%PEG 8000 for the sodium data set; and (5) 100 mM Mg-Acetate, 200 mM KCl, 50 mM LiCl, 50 mM Na-HEPES pH 7.0, 4% PEG 8000 for the potassium data set representing the substrate-free state.

### Structure determination

Diffraction data were collected at beamlines ID30A-1 and ID30B at ESRF (Grenoble, France) and processed with the XDS suite (Kabsch, 2010). The structures were solved by molecular replacement using Phaser in CCP4 (McCoy et al., 2007) and the RNA coordinates of PDB entry 4FAR and 4E8M (without solvent atoms) as the initial model (Marcia, 2016; Marcia et al., 2013a; Marcia and Pyle, 2012). The models were improved automatically in Phenix (Adams et al., 2010; Adams et al., 2004; Adams et al., 2002) and Refmac5 (Murshudov et al., 2011) and manually in Coot (Emsley, 2004), and finally evaluated by MolProbity (Davis, 2007). The figures depicting the structures were drawn using PyMOL Molecular Graphics System (Version 1.5.0.4, Schrödinger).

### pK_a_ calculations

Using MacroModel (v11.2, Schrödinger), we performed an OPLS4-based (Lu et al., 2021) conformational search of the input ligand structure and included in the next calculation the five most stable conformations of both ligand’s protonation states, for a total of 10 input conformations. Then, we employed the fully-analytic DFT-based protocol implemented in the Jaguar pKa computational tool with default parameters, allowing the use of zwitterionic functional groups (Bochevarov et al., 2016).

### Equilibrium molecular dynamics simulations

We used four structural models to perform MD simulations. Specifically, we modelled the holo-systems based on the structure of the intronistat B bound to the free intron (**Figure 2**, PDB id: 8OLS), and that of the ligand in complex with the 5’exon-bound intron (**Figure 4**, PDB id: 8OLZ). We modelled the corresponding apo-systems by removing the ligand from the holo-models, thus obtaining the free intron and 5’-exon-bound intron structures.

We used the AMBER force-field RNA.OL3 to parametrize the RNA molecules (Perez et al., 2007; Zgarbova et al., 2011). We used Joung and Cheatham (Joung and Cheatham, 2009), Panteva 12-6-4 non-bonded fixed-point charge (Panteva et al., 2015), and TIP3P models (Jorgensen et al., 1983) to parametrize ions, divalent ions and water, respectively.

We used the following protocol to set up and perform the production run for all the structural models. We performed Langevin dynamics simulations (Turq et al., 1977) with an integration time step of 2 fs, using AMBER (Salomon-Ferrer et al., 2013). We set the pressure at 1 atm using a Berendsen barostat with a relaxation time of 2 ps, while the temperature of 300K was controlled using a collision frequency γ = 1 per ps (Berendsen et al., 1984).

First, we performed energy minimization to relax the bulk water molecules, imposing 300 kcalꞏmolꞏÅ^2^ on the heavy atoms of the models, including the crystallized metals. Then, we smoothly thermalized the whole system at 300K using one NVT simulation of ∼1 ns, keeping the positional restraints used during the energy minimization. We then halved such restraints with a series of 3 ∼300-ps-long NVT simulations, performed two additional NPT simulations to relax the density of the systems to ∼1.01 gꞏcm^−3^, and further halved the restraints, which we removed during a third NPT run of ∼2 ns. We performed 2 independent simulation replicates for each structural model, resulting in 8 independent MD simulations and a total simulation time of 4 μs.

### Thermodynamic integration-based alchemical free energy calculations

Alchemical free energy calculations based on thermodynamic integration (TI) allow the estimation of the relative free energy of binding of two ligands, namely A and B, from molecular simulations that connect A and B bound or unbound states via a thermodynamic cycle (**Figure S8A**). This cycle is created by two thermodynamic paths that “alchemically” bridge the bound-A to bound-B states and unbound-A to unbound-B states, respectively, via a series of unphysical states. This is possible by introducing a coupling parameter λ that varies from zero (i.e., ligand A) to one (i.e., ligand B) and controls the unphysical decoupling of the contributions to the potential energy (i.e., van der Waals and Coulomb intra- and intermolecular interactions) of each ligand from the bound state and recouple them in the unbound state (water). In practice, two series of simulations are performed. One set of simulations alchemically transforms the ligand A into B when they are immersed in bulk water, thus representing the unbound state. The same is done for the bound state, thus transforming the ligand A into B when they are in complex with the receptor. Then, the ΔG associated with these alchemical transformations is estimated (ΔG_A_ and ΔG_B_, **Figure S8A**), and, building upon the thermodynamic cycle in **Figure S8A**, it is possible to calculate ΔΔG_AB_ = ΔG_B_-ΔG_A_. A detailed description of the computational method and its theoretical framework is out of the scope of this study and can be found elsewhere (Cournia et al., 2017).

So far, alchemical free energy calculations have been reliably used to guide the structure-based design of small molecule drugs and predict the effect of functional groups on the potency of the newly designed compound before it is synthesized and tested (De Vivo et al., 2016; Jorgensen, 2009). However, for alchemical free energy calculations to be effectively reliable, the binding mode of the ligands must be modeled very accurately to account for all the interactions responsible for the experimentally observed binding potency. Otherwise, the free energy estimates of the bound state could be dramatically affected, resulting in an inaccurate estimate of the total ΔΔG_AB_ (Hahn et al., 2022; Mey et al., 2020). Building upon this evidence, in this study we decided to use alchemical free energy calculations in a reverse approach than usually done. That is, instead of relying on a validated binding mode to estimate compounds’ ΔΔG_AB_ and computationally drive the improvement of binding potency, here we relied on experimentally computed ΔΔG_AB_ of small molecules to compare with our estimates so as to computationally validate the ligands’ binding mode.

We employed the GPU-TI code implemented in AMBER22 to perform alchemical free energy calculations (Lee et al., 2017; Salomon-Ferrer et al., 2013). The ligand-bound structural models were prepared as follows. Given a couple of ligands A and B, where B is bigger than A, then the binding of ligand A was modeled at the active site of the intron building upon the pose of intronistat B. This has been possible since we evaluated ΔΔG only of ligands having the same scaffold as intronistat B (i.e., the pyrogallol moiety and the benzofuran core). The ligand-A bound complex was then equilibrated via classical MD simulations using the same protocol mentioned in the previous paragraph, and a production run of 100 ns was performed to properly relax the system (**Figure S8B**). The last frame’s coordinates obtained via this MD simulations were used to set up the topology for the TI-based alchemical transformations, which were performed disabling the SHAKE algorithm, setting the time step to 1 fs and the collision frequency to 5, as well as using the Monte Carlo barostat with a pressure relaxation time of 1.0 ps to keep the pressure at 1.01325 bar. TI simulations followed a λ-scheme, similar to previous works (**Figure S8B**, (He et al., 2020)), starting with a preliminary thermalization (1 ns) and pressurization (1.5 ns) of the system in the NVT and NPT ensemble, respectively, setting λ=0.5. At this point, a divergent equilibration scheme is initiated to equilibrate all twelve λ-windows (λ: 0.00922, 0.04794, 0.11505, 0.20634, 0.31608, 0.43738, 0.56262, 0.68392, 0.79366, 0.88495, 0.95206, and 0.99078 and weighs 0.02359, 0.05347, 0.08004, 0.10158, 0.11675, 0.12457) via two parallel branches of NPT simulations (2.5 ns, each λ-window). One branch leads to the progressive equilibration of the windows from λ = 0.5 to λ = 0.00922, the other from λ = 0.5 to λ = 0.99078, such that each simulation is started from an equilibrated frame taken from the previous λ-window. Finally, production simulations are performed using Hamiltonian replica exchange via λ-hopping, with an exchange rate of 10 ps, for a total simulation time of 5 ns per λ-window.

For the ligand-in-water structural models, the compounds were immersed in a box of TIP3P water whose edges were separated by 20 Å from the solute, and monovalent ions were added to neutralize the system when needed. We used the same simulation protocol described for the ligand-bound state. For both the bound and unbound ligands’ states, in case of non-neutral transformations (i.e., the net charge of one compound differs from the one of the other), we applied the so-called co-ion approach (Chen and Roux, 2015; Wallace and Shen, 2012). Here, one metal ion with appropriate charge is transformed into one water molecule to balance the net charge of the system during the compounds’ transformation. Smoothstep softcore potentials were used to treat ligands’ non-common atoms (Lee et al., 2020).

Overall, we performed the following TI-based alchemical free energy calculations, cumulating ∼1μs of simulation time: *i)* compound 8 → intronistat B as bound to the free intron (**Figures S14-15**, 3 replicas); *ii)* compound 8 → intronistat B as bound to the intron in complex with the 5’exon (**Figures S9-10**, 3 replicas); *iii)* compound 8 → compound 12 as bound to the intron in complex with the 5’exon (**Figures S11A-12A**, 1 simulation); *iv)* compound 12 → compound 17 as bound to the intron in complex with the 5’exon (**Figures S11B-12B**, 1 simulation); *v)* compound 17 → intronistat A as bound to the intron in complex with the 5’exon (**Figures S11C-12C**, 1 simulation).

### Chemistry

#### General considerations

All the commercially-available reagents and solvents were used as purchased from vendors without further purification. Dry solvents were purchased from Sigma-Aldrich. Intronistat B hydrobromide (≥98%) was purchased from Sigma-Aldrich and used for the assays reported in the present work. A separate batch of intronistat B hydrobromide was synthesized as described in WO2019147894, and used as a starting material for the synthesis of ARN25850 [2-(2,6-dibromo-3,4,5-trihydroxybenzoyl)-*N*-(2-(pyrrolidin-1-yl)ethyl)benzofuran-5-carboxamide hydrobromide, **Scheme S1**]. NMR data were collected on a Bruker Avance III 400 MHz (^1^H) and 100 MHz (^13^C). Spectra were acquired at 300 K, using deuterated dimethylsulfoxide (DMSO–*d_6_*). For ^1^H-NMR, data are reported as follows: chemical shift, multiplicity (s= singlet, d= doublet, dd= double of doublets, t= triplet, q= quartet, m= multiplet), coupling constants (Hz) and integration. UPLC/MS analyses were run on a Waters ACQUITY UPLC/MS system consisting of an SQD (Single Quadrupole Detector) Mass Spectrometer equipped with an Electrospray Ionization interface and a Photodiode Array Detector. PDA range was 210-400 nm. The mobile phase was 10mM NH_4_OAc in H_2_O at pH 5 adjusted with AcOH (A) and 10mM NH_4_OAc in CH_3_CN-H_2_O (95:5) at pH 5 (B). The analysis was performed on an ACQUITY UPLC BEH C18 (100×2.1 mmID, particle size 1.7 μm) with a VanGuard BEH C18 pre-column (5×2.1 mmID, particle size 1.7 µm) and the mobile-phase B proportion increased from 10 % to 90 % in 6 min. Electrospray ionization in positive and negative mode was applied in the mass scan range 100-650Da.

#### Synthesis of di-brominated intronistat B derivative, ARN25850

1,3-Dibromo-5,5-dimethylhydantoin (DBDMH) (48.5 mg, 0.17 mmol) was added portionwise to a suspension of intronistat B hydrobromide (60.0 mg, 0.12 mmol) in CHCl_3_ dry (3.5 mL) at room temperature under argon. The resulting suspension was stirred for 72 hours; the limpid supernatant was decanted from the solid material. This latter was triturated with isopropanol (1 mL), followed by 10% isopropanol in chloroform (twice), and dichloromethane (once). The resulting solid was dried *in vacuo* (60°C) to provide the title compound as a brownish solid (16 mg, 20%). UPLC/MS: Rt: 2.50 min. MS (ESI) m/z: 566.97 [M-H]^-^. [M-H]^-^ calculated for C_22_H_20_Br_2_N_2_O_6_ = 566.96, (**Figure S16**). ^1^H (400 MHz, DMSO-*d_6_*): δ 9.70 (bs, 2H), 9.50 (bs, 1H), 8.86 (t, *J* = 5.2 Hz, 1H), 8.35 (d, *J* = 1.8 Hz, 1H), 8.09 (dd, *J* = 8.9, 1.8 Hz, 1H), 7.87 (d, *J* = 8.8 Hz, 1H), 7.70 (s, 1H), 3.64 (m, 4H), 3.08 (m, 2H), 2.02 (m, 2H), 1.87 (m, 2H). (One CH_2_ overlaps with water, signal recovered by HSQC, 3.37ppm). ^13^C NMR (101 MHz, DMSO-*d_6_*) δ 183.0 (Cq), 166.4 (Cq), 157.0 (Cq), 151.9 (Cq), 143.6 (Cq, 2C), 137.0 (Cq), 130.5 (Cq), 129.8 (Cq), 128.4 (CH), 126.8 (Cq), 124.0 (CH), 117.7 (CH), 112.4 (CH), 98.7 (Cq, 2C), 53.6 (CH_2_, 2C), 53.3 (CH_2_), 36.0 (CH_2_), 22.5 (CH_2_, 2C). ^1^H, ^13^C, COSY and HSQC NMR spectra and MS and VIS-UV spectra are represented in **Figures S16-S20**.

## SUPPLEMENTAL MATERIAL

### Supplemental Figures

**Figure S1.**
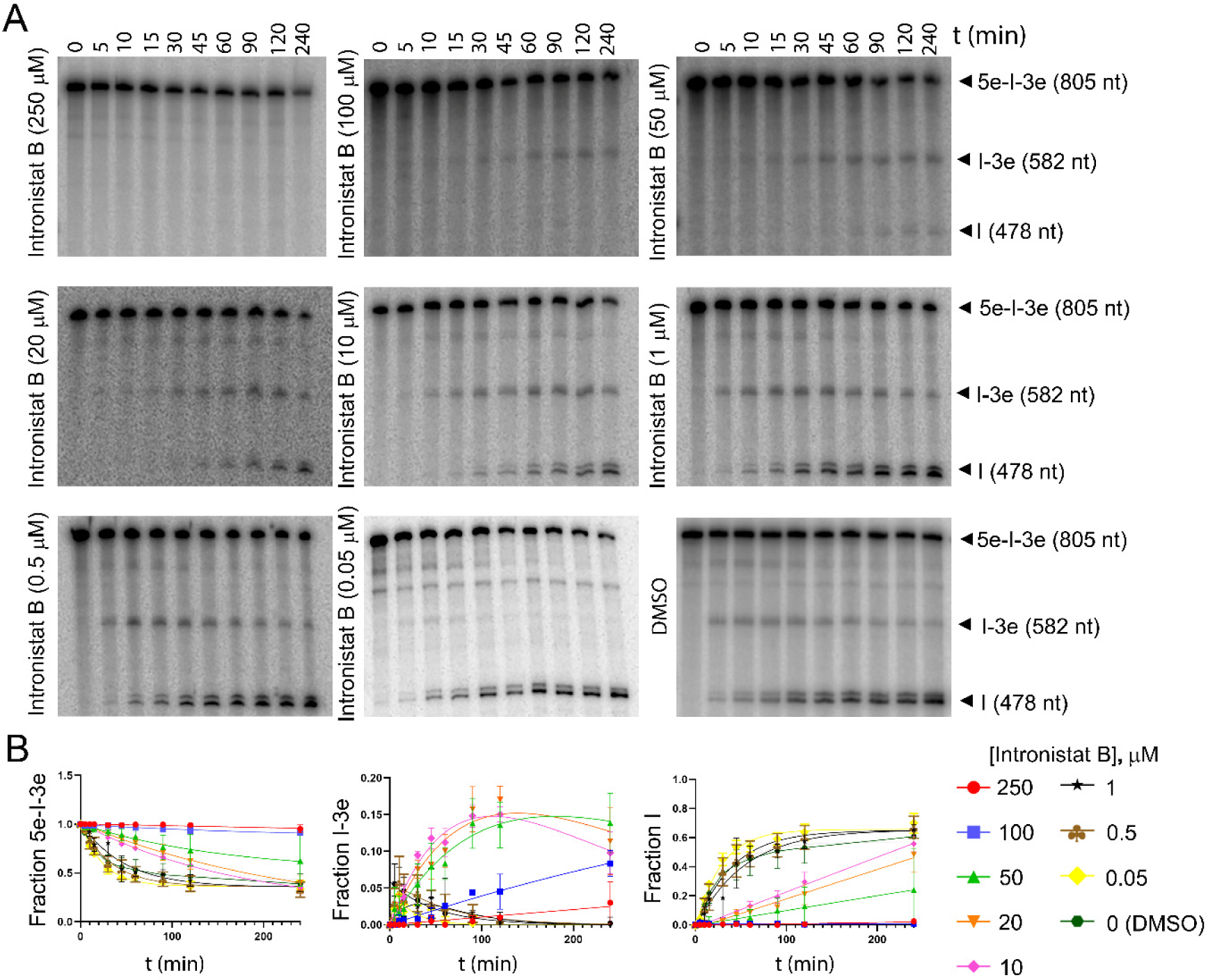
Kinetics of splicing inhibition. **(A)** Splicing kinetics in the presence of Intronistat B at all concentrations tested in this study. The relative rate constants are listed in **Table S1**. Precursors are indicated as 5e-I-3e (nt length in parenthesis). Intermediate (I-3e) and linear intron (I) migrate as double bands because of cryptic cleavage sites, as explained previously (Manigrasso et al., 2020; Marcia and Pyle, 2012). **(B)** Evolution of the populations of precursor (5e-I-3e, left panel), intermediate (I-3e, middle panel), and linear intron (I, right panel) over time. Error bars represent standard errors of the mean (s.e.m.) calculated from n = 3 independent experiments.

**Figure S2.**
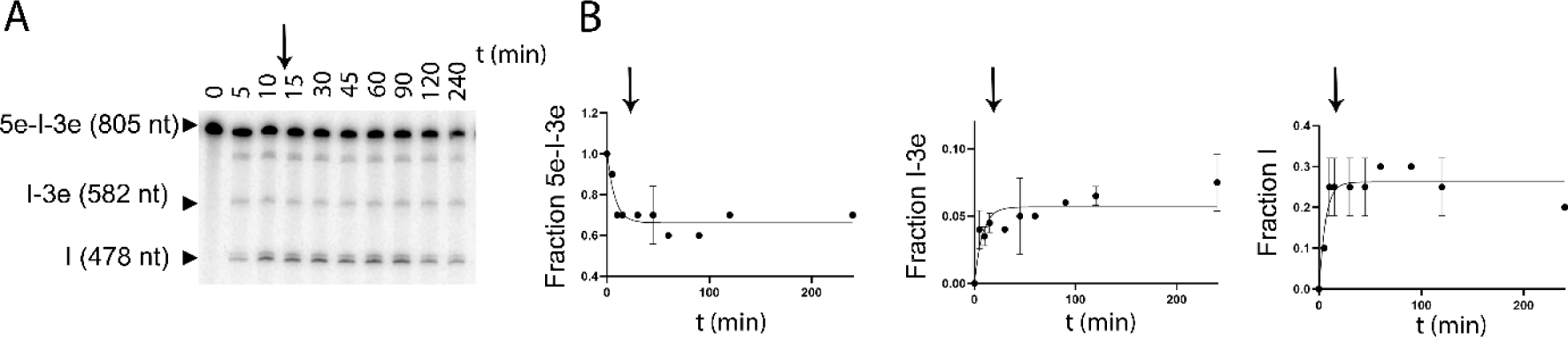
Intronistat B targets folded, active group II introns. **(A)** Representative splicing kinetics in the presence of 250 μM intronistat B, which is added 10 min after the start of the splicing reaction (black arrow). **(B)** Evolution of the populations of precursor (5e-I-3e, left panel), intermediate (I-3e, middle panel), and linear intron (I, right panel) over time. The black arrows indicate the time when intronistat B was added to the reaction.

**Figure S3.**
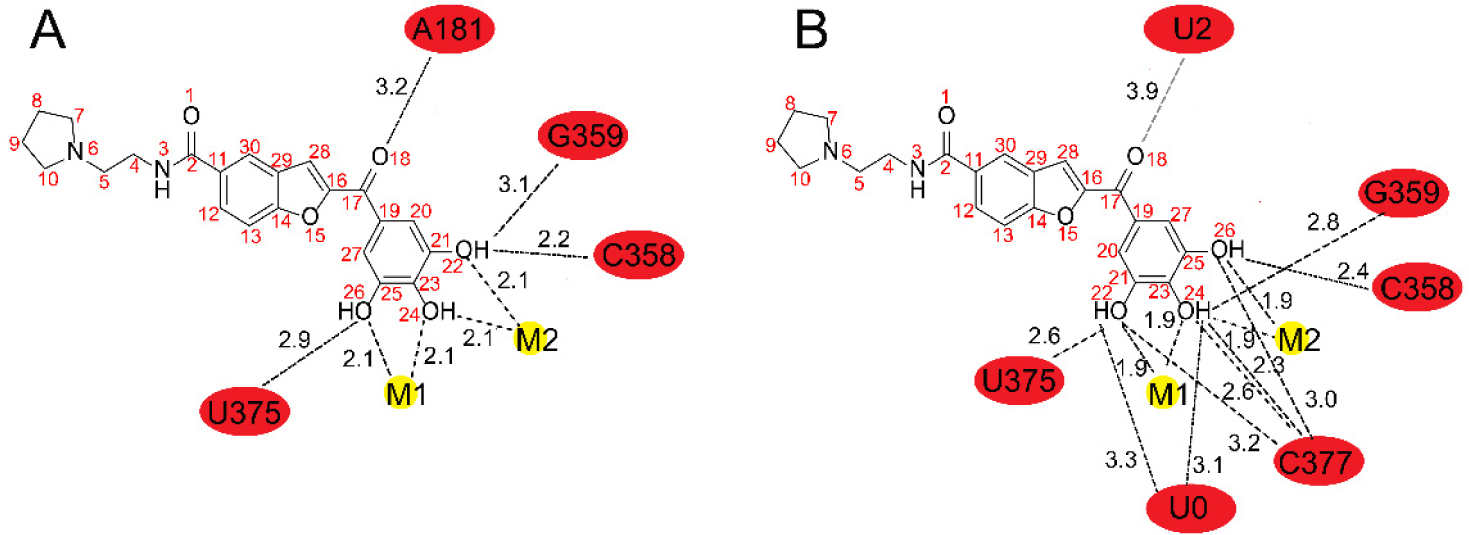
Intronistat B chemical structure and interactions with the active site of group II introns. **(A)** Intronistat B and its interactions with the group II intron in the exon free state (related to Figure 2). **(B) I**ntronistat B and its nteractions with the group II intron in the exon bound state (related to Figure 4). Black dotted lines indicate the interactions between the atoms of intronistat B and the group II intron active site nucleotides (red circles) or catalytic metals (yellow circles). The weak contact between intronistat B O18 and the U2 O4’ atom is indicated by a gray dotted ine. Distances are indicated in Å next to each dotted line.

**Figure S4.**
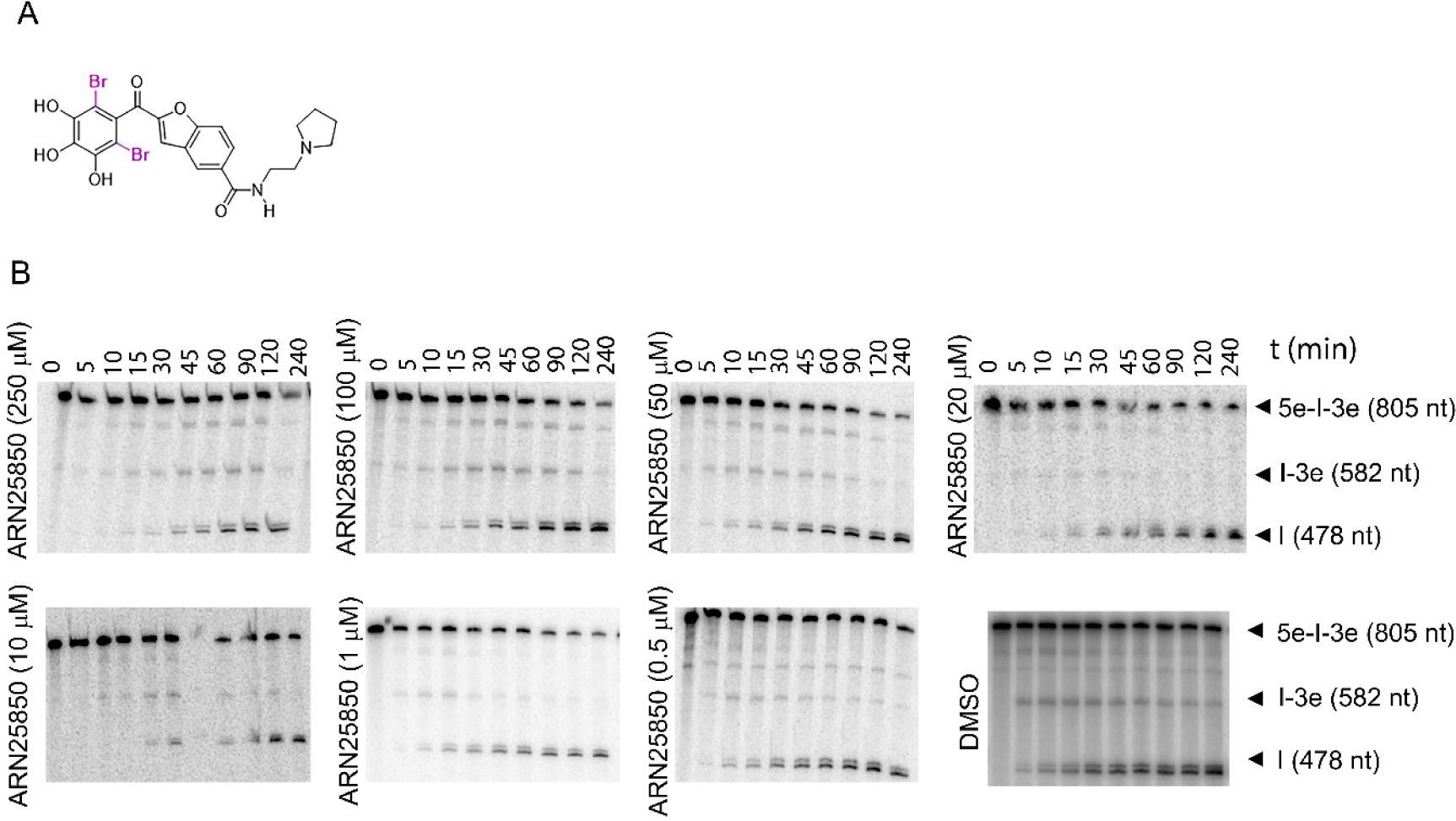
Group II intron inhibition by the di-brominated intronistat B derivative, ARN25850. **(A)** Chemical structure of ARN25850. **(B)** Representative splicing kinetics in the presence of different concentrations of ARN25850. The relative rate constants are listed in **Table S1**.

**Figure S5.**
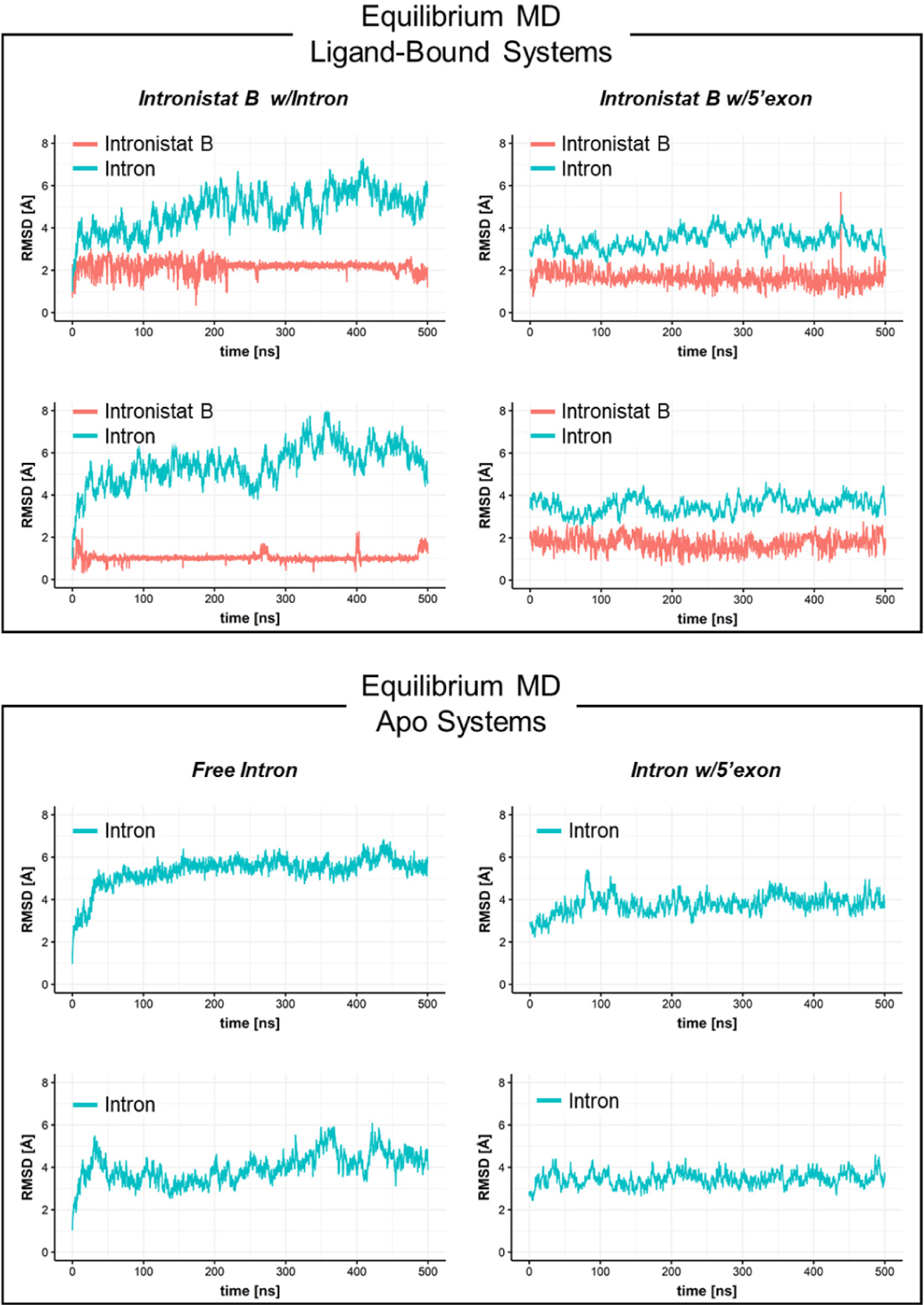
MD simulations of the free and 5’exon-bound intron. On the top four panels, the RMSD value of the intron (cyan) and the intronistat B (red) is reported as a function of the simulation time for the free (left) and 5’exon-bound (right) system. On the bottom four panels, the RMSD of the free (left) and 5’exon-bound (right) intron is reported.

**Figure S6.**
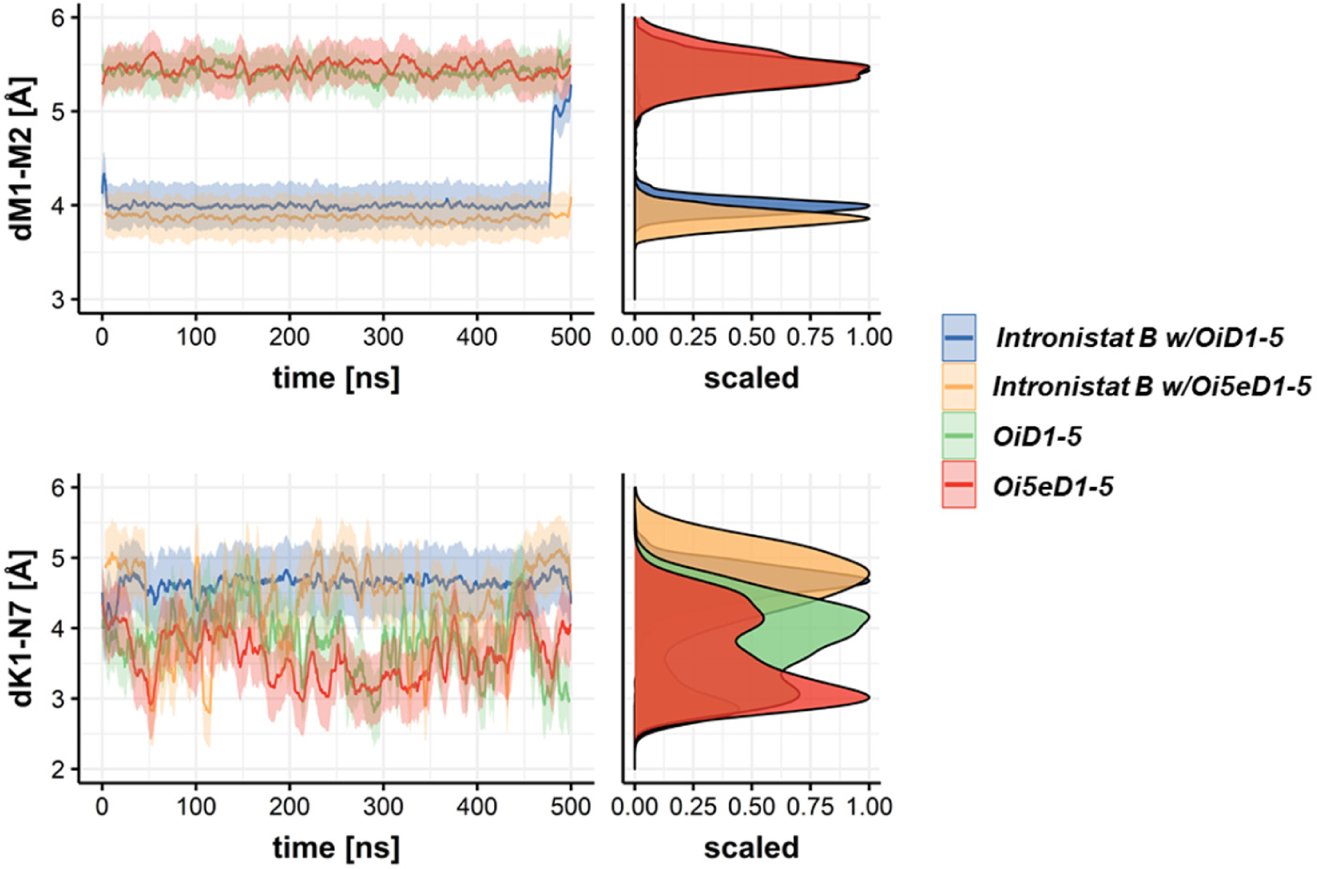
Binding of splicing modulators alters the functional dynamics of intron’s catalytic features. The Figure reports MD replicas in support of simulations shown in Figure 3.

**Figure S7.**
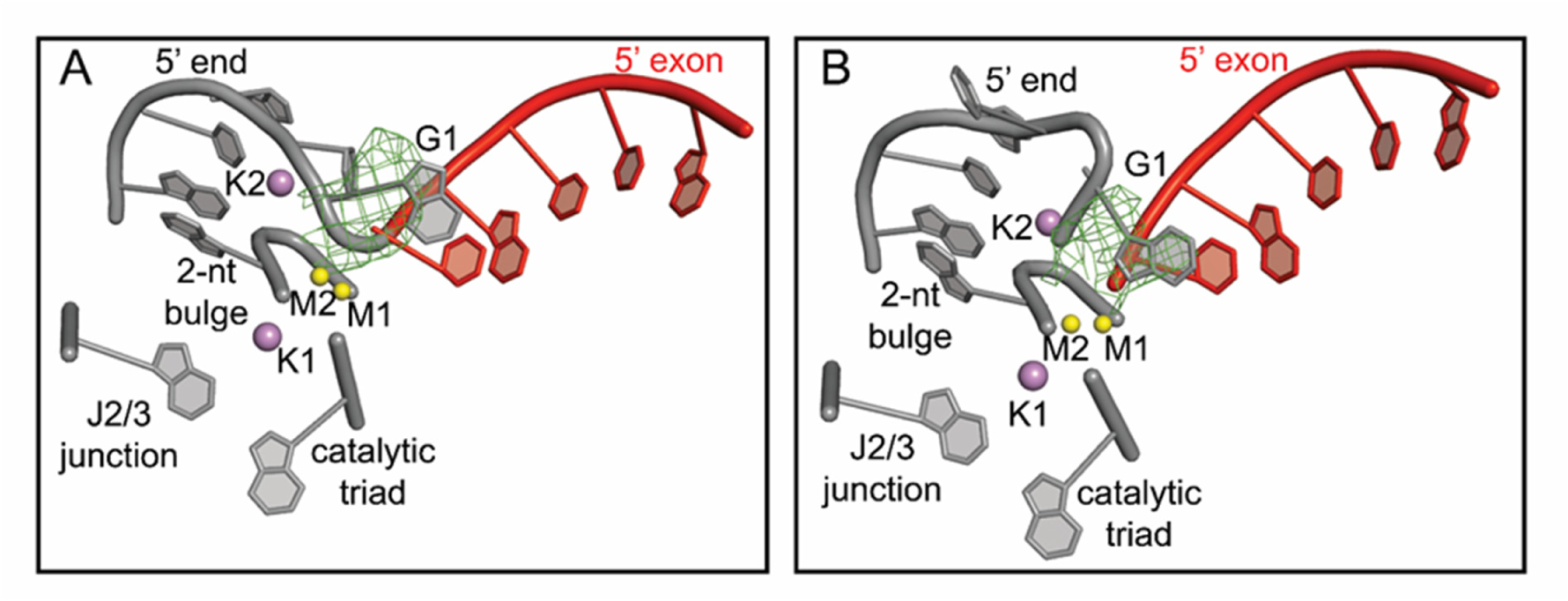
The 5’-splice junction outcompetes intronistat B in the active site. **(A)** Crystal structure of Oi5eD1-5 in the pre-catalytic stage in the presence of Ca_2+_ (yellow spheres), K_+_ (purple spheres) and intronistat B. **(B)** Crystal structure of Oi5eD1-5 in the post-catalytic stage in the presence of Mg_2+_ (yellow spheres), K_+_ (purple spheres) and intronistat B. The F_o_−F_c_ electron density omit map calculated by omitting intron residue G1 and contoured at 3σ is represented as a green mesh in both panels. The 5’-exon is represented as red sticks in both panels. Intronistat B is not bound to the active site under these conditions.

**Figure S8.**
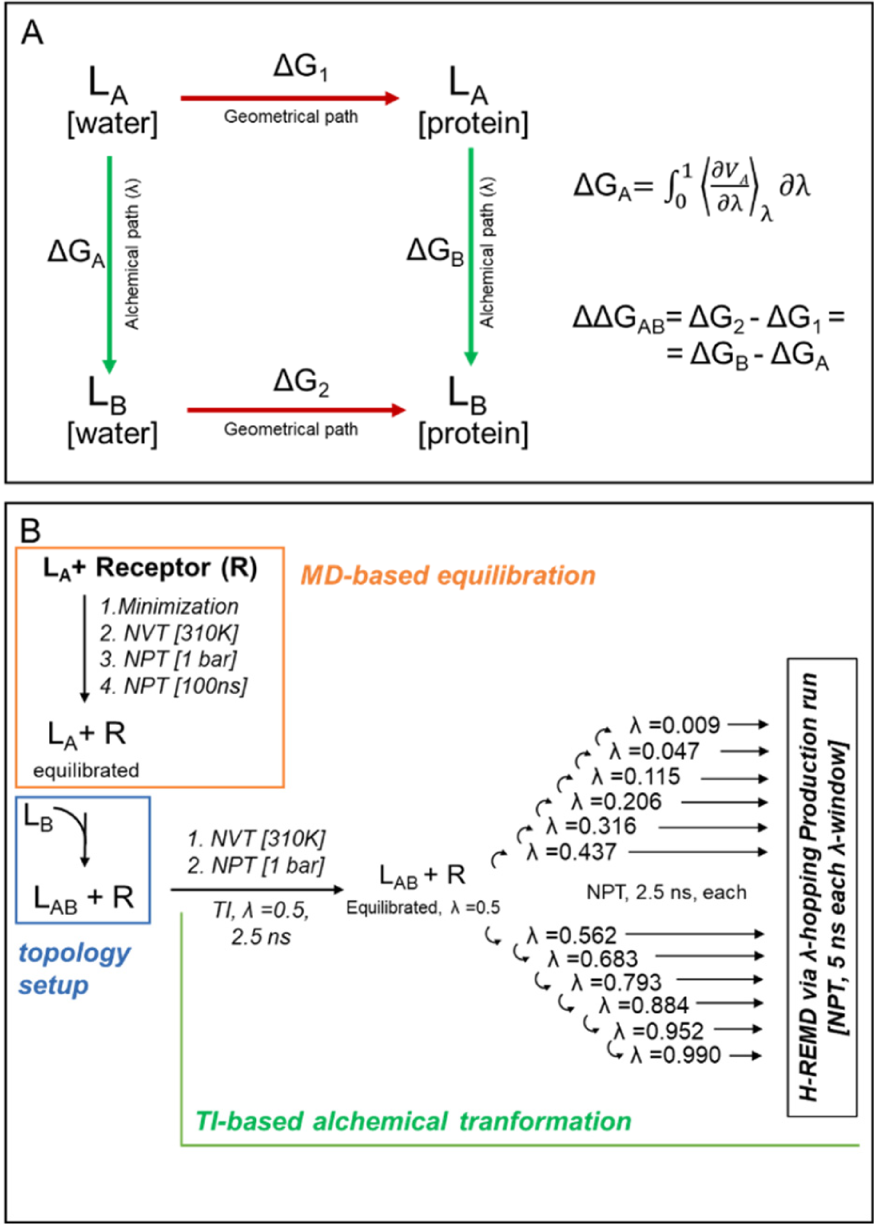
Thermodynamic integration-based alchemical free energy calculations. **(A)** The thermodynamic cycle at the basis of the estimations of relative binding free energy. Additionally, the equation for deriving the ΔG along every alchemical path, as well as that for deriving the ΔΔG is reported. **(B)** The protocol for the alchemical free energy calculations.

**Figure S9.**
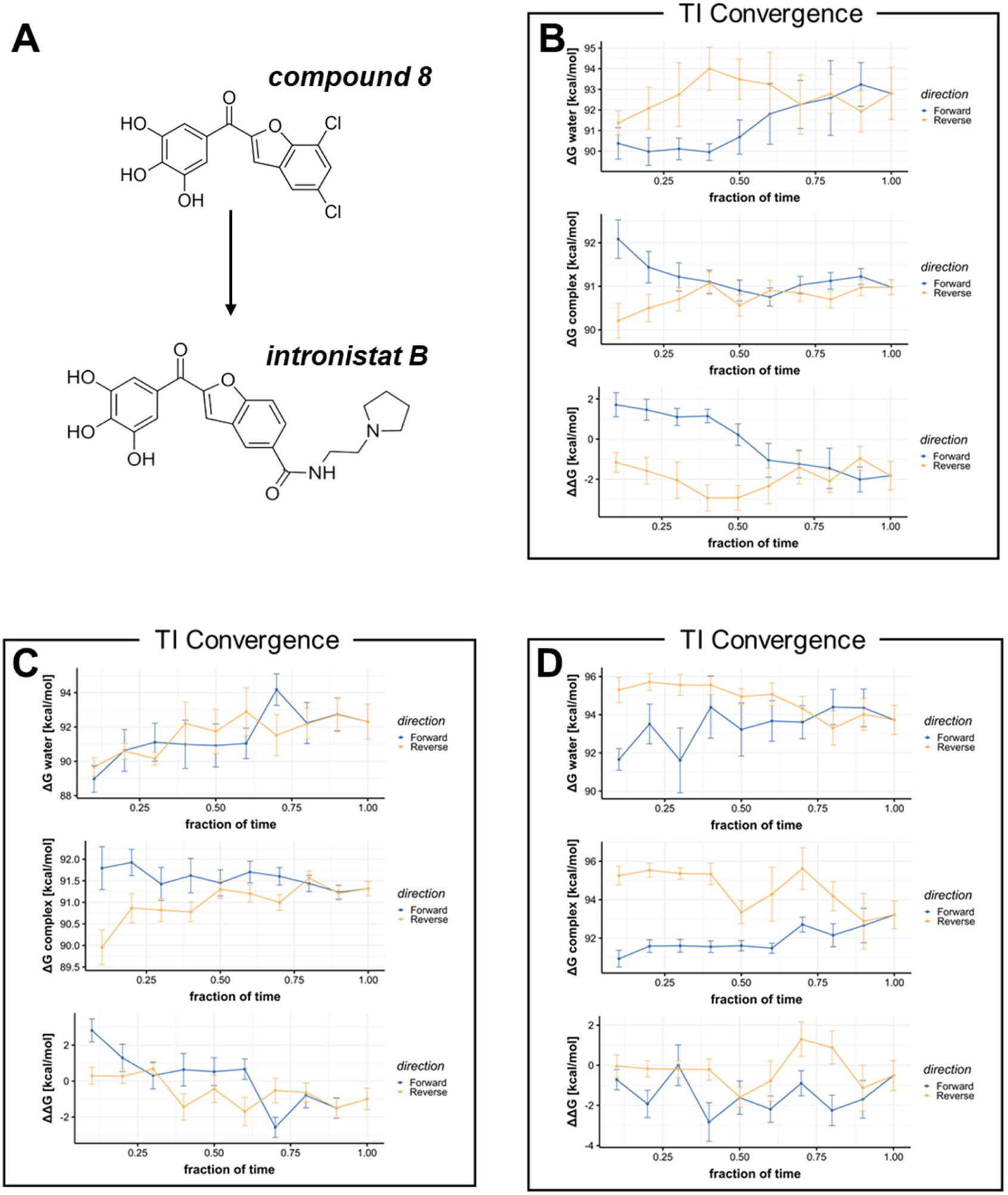
Alchemical free energy calculations for compound 8 and intronistat B bound to the intron in complex with the 5’-exon. **A**) The 2D structures of the compounds are reported. The convergence of the forward (blue) and reverse (yellow) estimates of the ΔG of the ligands in water and as bound to the receptor, as well and their ΔΔG is reported of each of three simulations replica (**B-D**).

**Figure S10.**
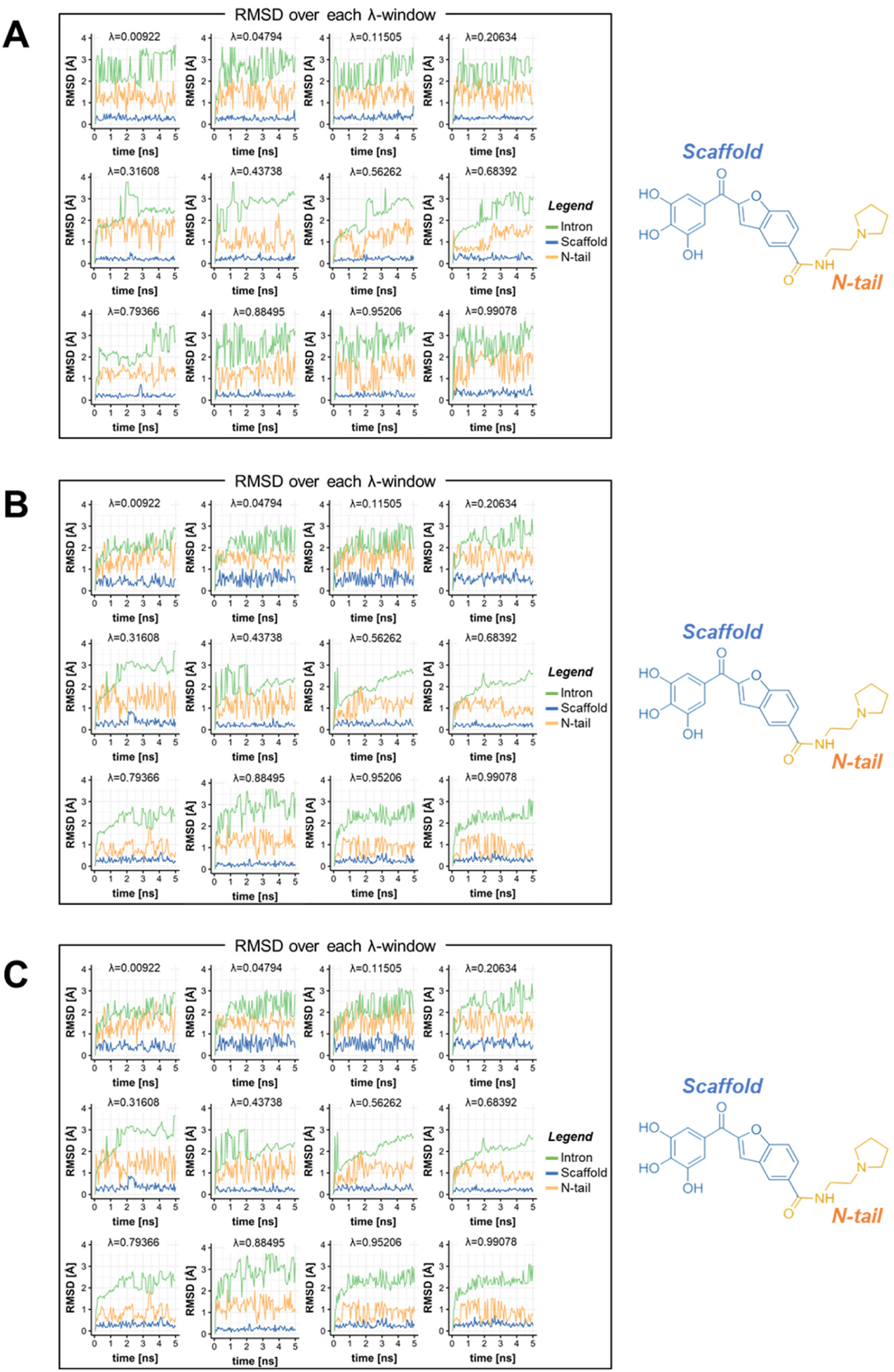
Ligands binding stability during alchemical free energy calculations for compound 8 and intronistat B bound to the intron in complex with the 5’-exon. **A-C)** The RMSD values of the intron (green), the intronistat B benzofuran scaffold (blue), and its N-tail (yellow, coloring scheme following that of Figure 5), are reported as a function of simulation time at each lambda window, for three simulations replicate. High flexibility is shown by the N-tail but not by the benzofuran scaffold (RMSD<0.5Å) in all windows. This results in a better convergence of the ΔΔG estimates.

**Figure S11.**
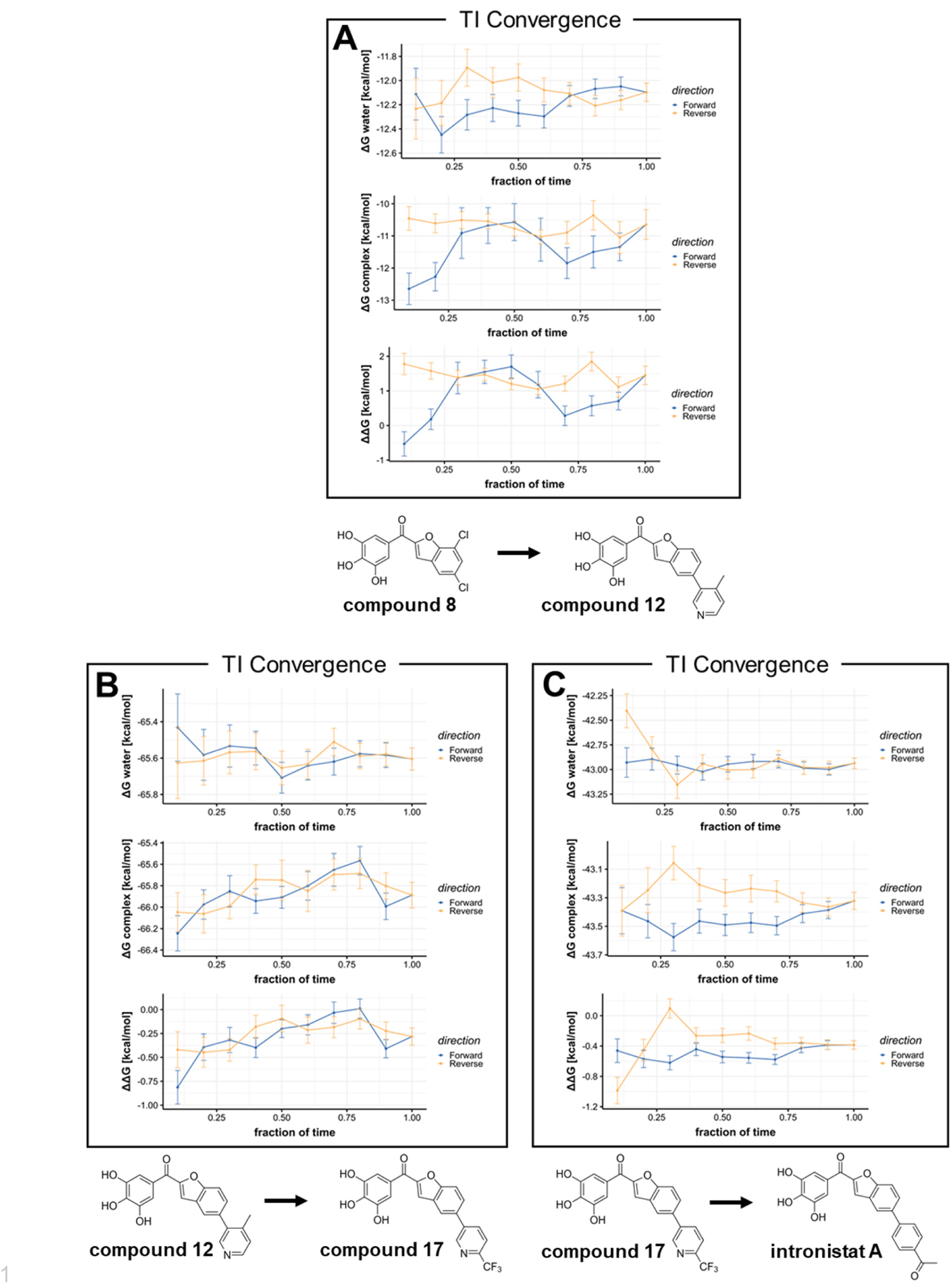
Alchemical free energy calculations for additional intronistat B analogues as bound to the intron in complex with the 5’-exon. The convergence of the forward (blue) and reverse (yellow) estimates of the ΔG of the ligands in water, as bound to the receptor and their ΔΔG is reported for the alchemical transformation of compound 8 and compound 12 (**A**), compound 12 and compound 17 (**B**), as well as compound 17 and intronistat A (**C**). The 2D structure of the compounds involved in the alchemical transformation is reported below each panel.

**Figure S12.**
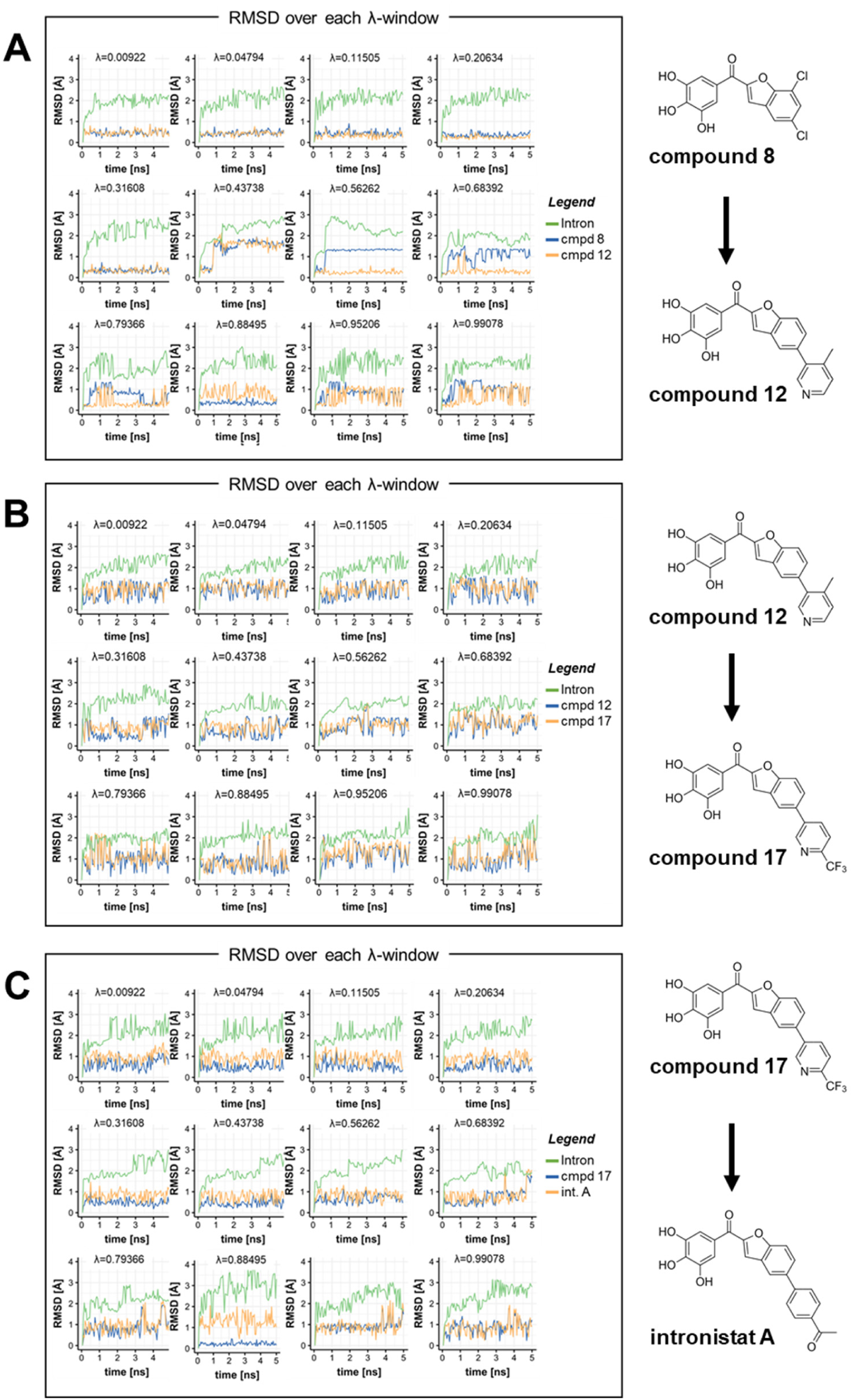
Ligands binding stability during alchemical free energy calculations of intronistat B analogues as bound to the intron in complex with the 5’-exon. **A-C)** The RMSD values of the intron (green), the intronistat B benzofuran scaffold (blue), and its N-tail (yellow, coloring scheme following that of Figure 5), are reported as a function of simulation time at each lambda window, for the alchemical transformation of compound 8 and compound 12 (**A**), compound 12 and compound 17 (**B**), as well as of compound 17 and intronistat A (**C**). The 2D structure of the compounds involved in the alchemical transformation is reported beside each panel.

**Figure S13.**
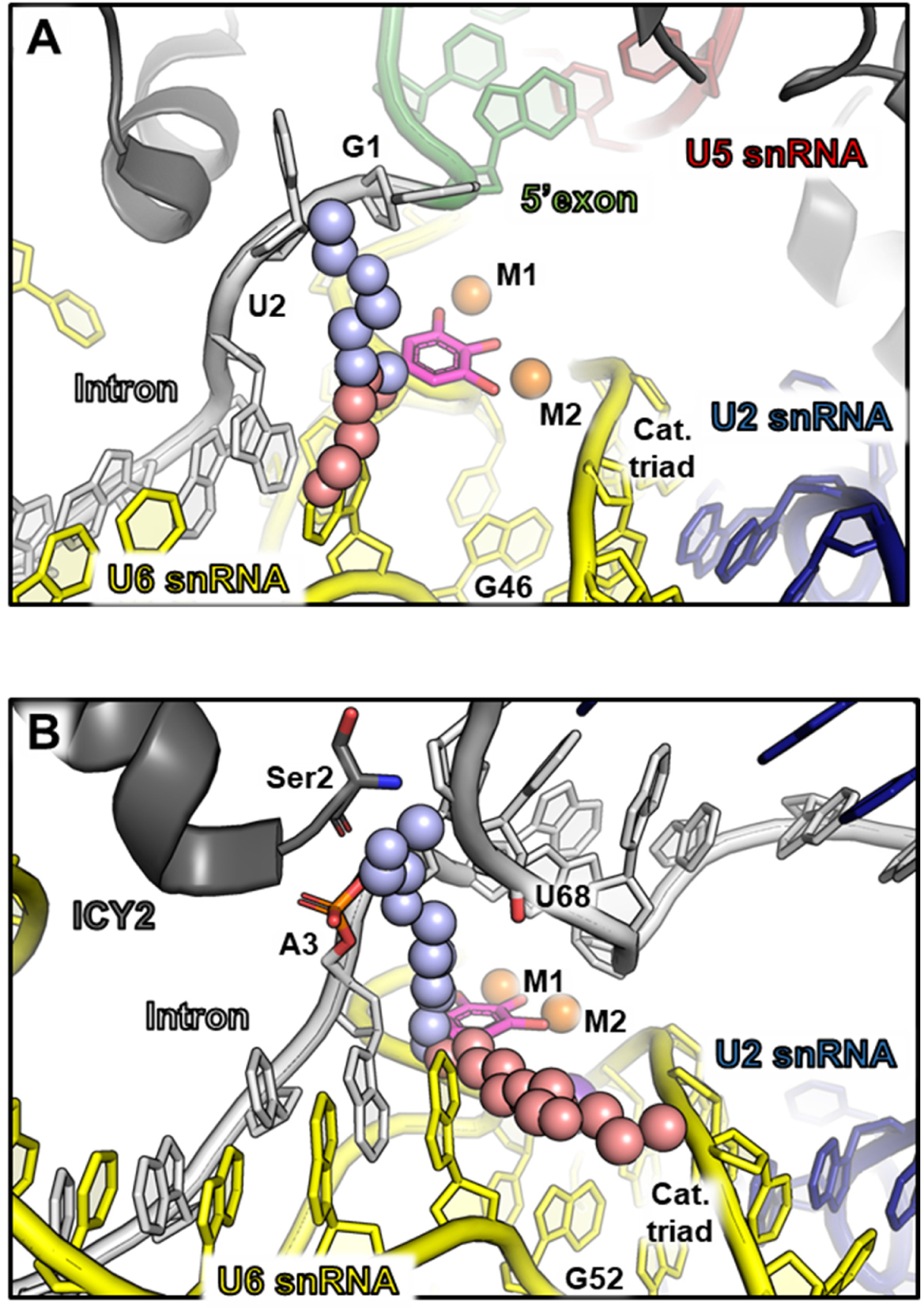
Putative binding modes of intronistat B analogues at the active site of the spliceosome at different stages of catalysis. **A)** Intronistat B derivatives bound at catalytic core of the human B* spliceosomal complex (PDB id: 5Z58). **B**) A similar binding pose can be modeled for intronistat B derivatives when bound to the Ci spliceosomal complex from *S. cerevisiae* (PDB id: 7B9V). Notably, in both cases, a two-metal-ion binding compound similar to intronistat B would locate in proximity of the splice junctions or the intron nucleotides, suggesting that sequence-specific contacts between small molecules and the spliceosomal complex can be possibly engaged upon having anchored its structurally-conserved active site.

**Figure S14.**
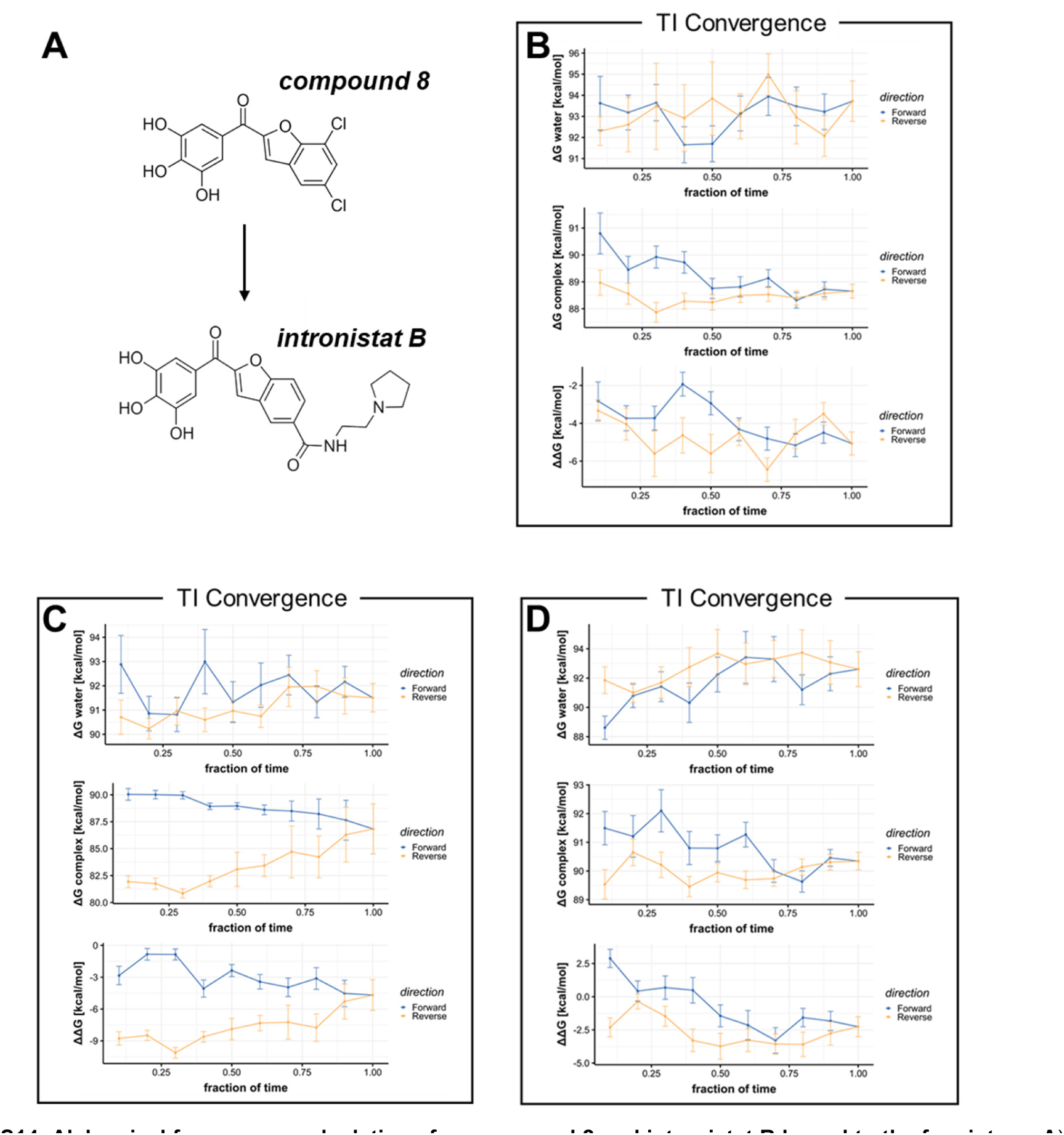
Alchemical free energy calculations for compound 8 and intronistat B bound to the free intron. **A)** The 2D structures of the compounds are reported. The convergence of the forward (blue) and reverse (yellow) estimates of the ΔG of the ligands in water and as bound to the receptor, as well their ΔΔG is reported of each of three simulations replica (**B-D**).

**Figure S15.**
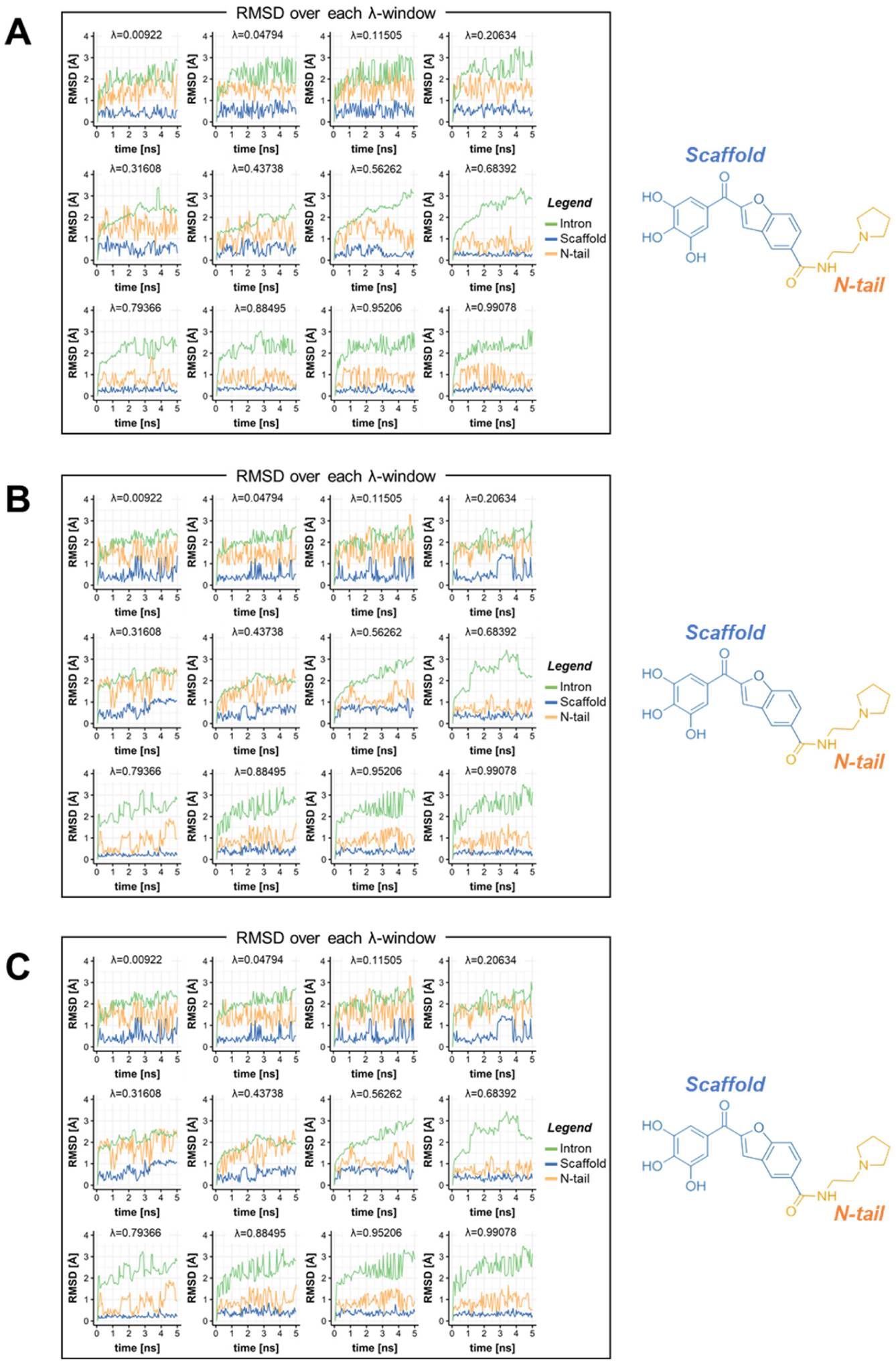
Ligands binding stability during alchemical free energy calculations for compound 8 and intronistat B bound to the free intron. **A-C)** The RMSD values of the intron (green), the intronistat B benzofuran scaffold (blue), and its N-tail (yellow, coloring scheme following that of Figure 5), are reported as a function of simulation time at each lambda window, for the three simulations replicate. High flexibility is shown by the N-tail and the benzofuran scaffold in several windows, as highlighted by their RMSD fluctuations greater than 2Å and 1Å, respectively. This results in the poor convergence of the ΔΔG estimates.

**Figure S16.**
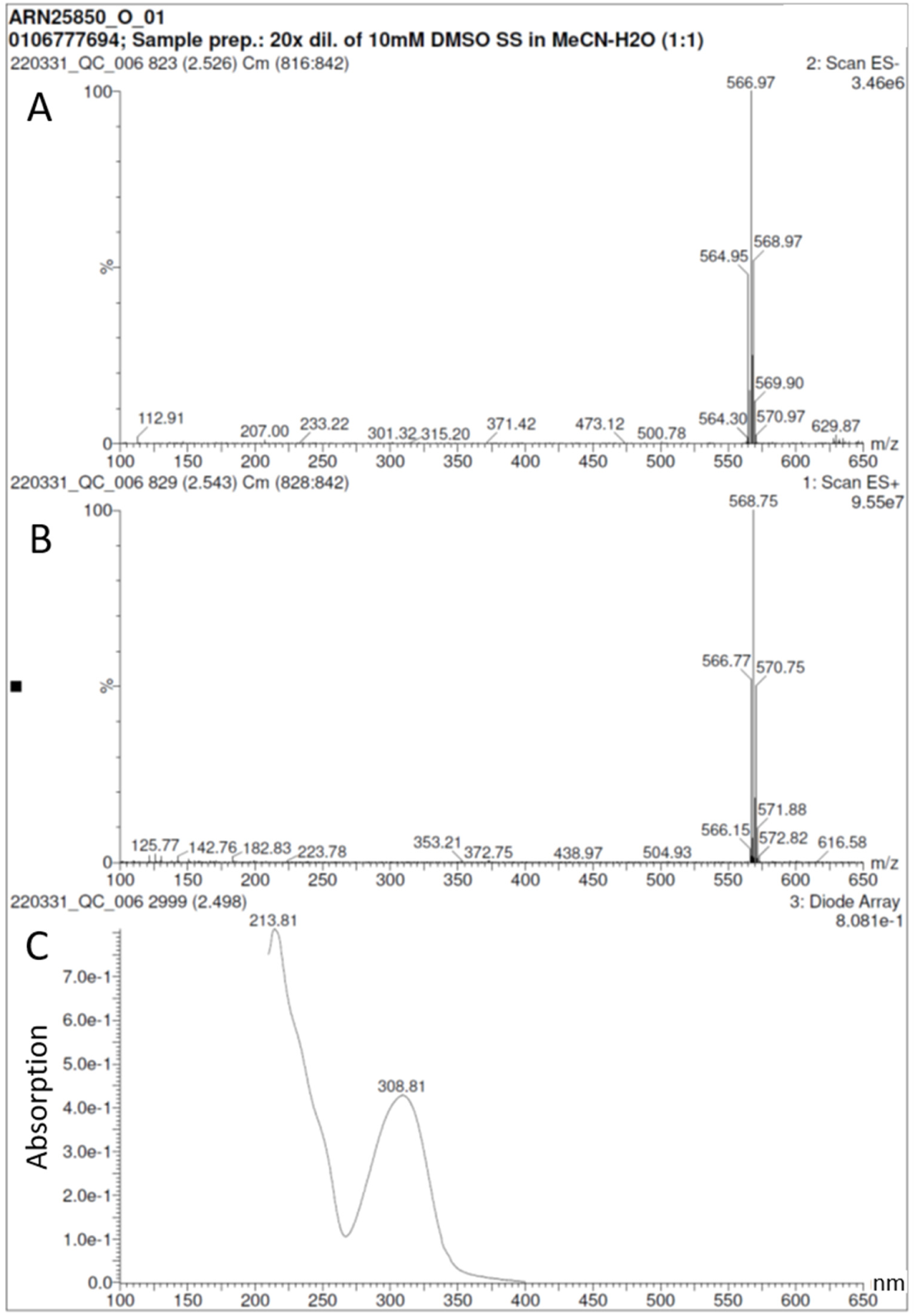
Synthesis of dibromo-intronistat B hydrobromide, ARN25850. UPLC-MS and UV spectra.

**Figure S17.**
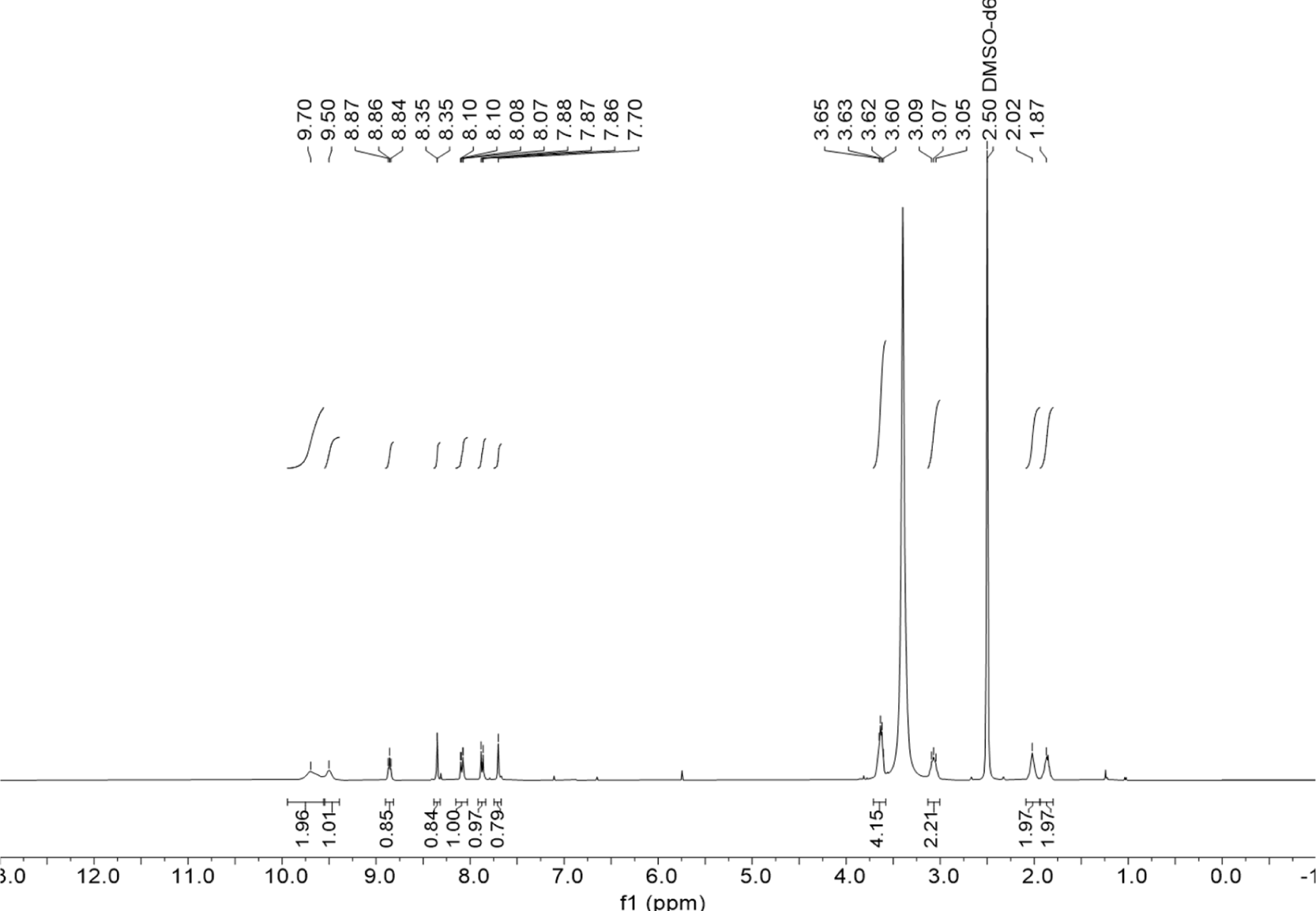
Synthesis of dibromo-intronistat B hydrobromide, ARN25850. ^1^H-NMR spectrum (DMSO-*d_6_*, 400MHz).

**Figure S18.**
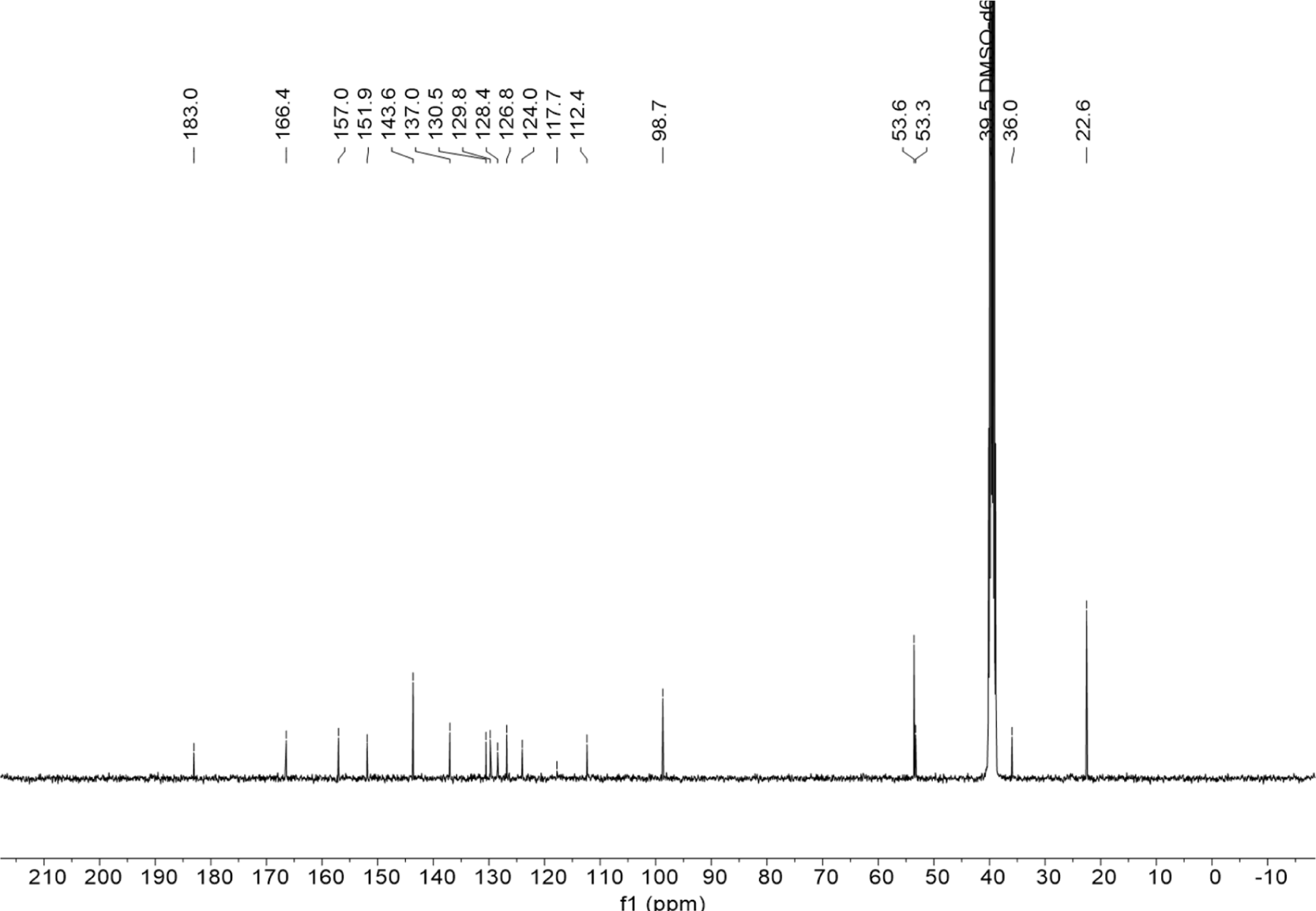
Synthesis of dibromo-intronistat B hydrobromide, ARN25850. ^13^C-NMR spectrum (DMSO-*d_6_*, 400MHz).

**Figure S19.**
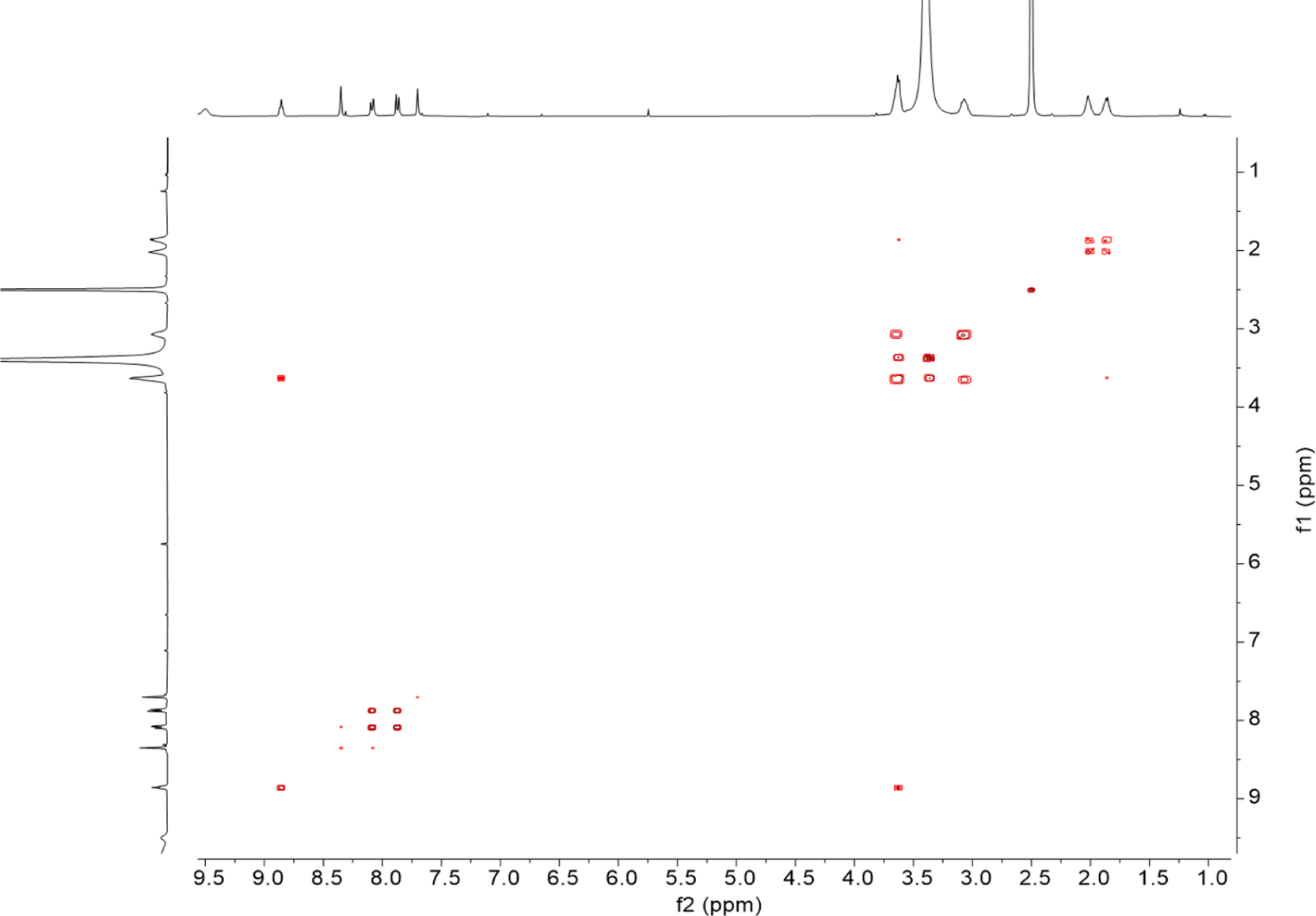
Synthesis of dibromo-intronistat B hydrobromide, ARN25850. COSY spectrum (DMSO-*d_6_*, 400MHz).

**Figure S20.**
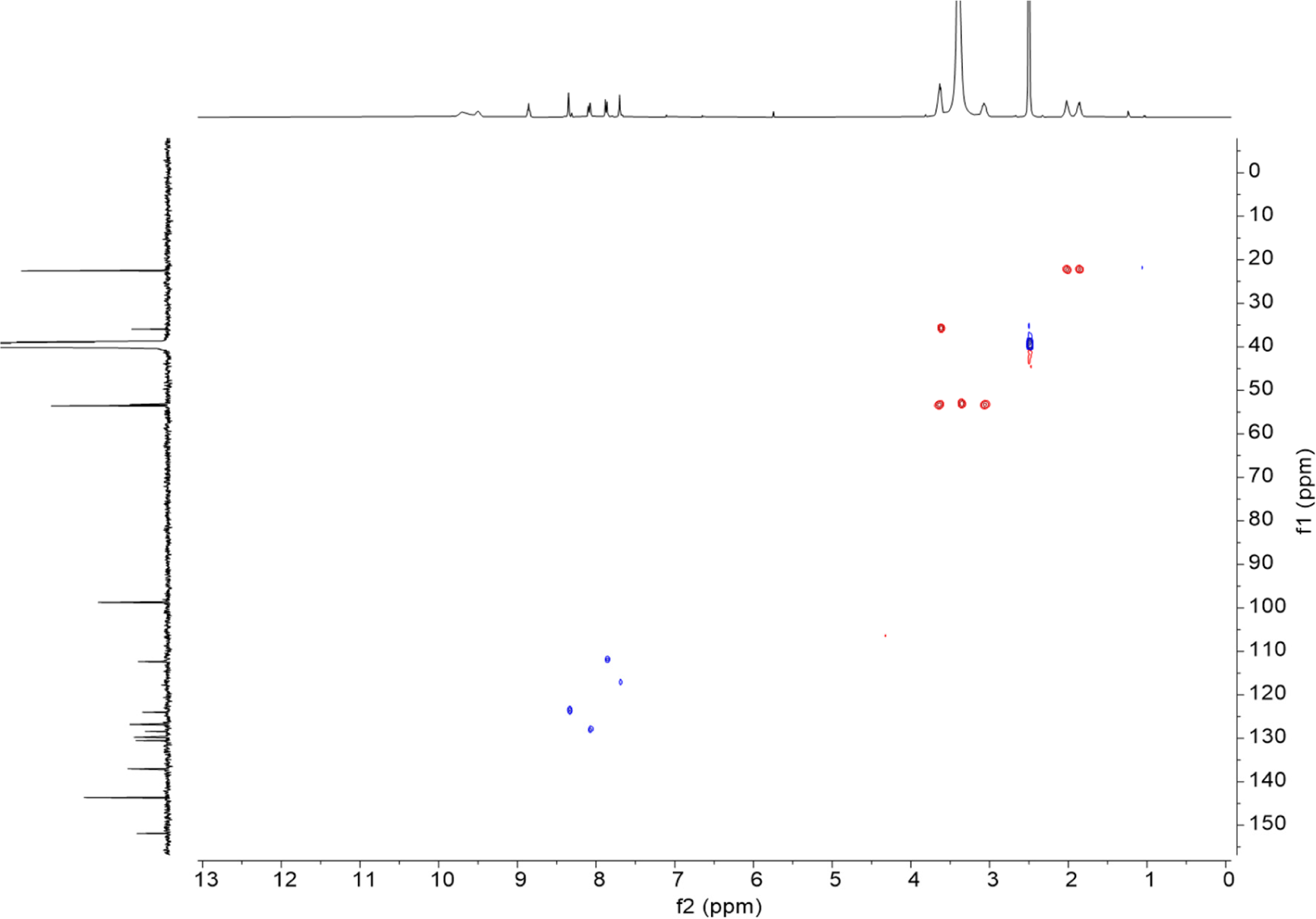
Synthesis of dibromo-intronistat B hydrobromide, ARN25850. HSQC spectrum (DMSO-*d_6_*, 400MHz).

### Supplemental Tables

**Table S1:** Kinetics rate constants. Splicing rate constants of the first (k_1_) and second (k_2_) steps of splicing of *O. iheyensis* group IIC intron in the presence of all concentrations of intronistat B and the di-brominated intronistat B derivative tested in this study. The total rate constant (k = k_1_ + k_2_) is also reported for easier comparison with previously reported rate constants of the mitochondrial ai5γ intron (Fedorova et al., 2018). N.D. = not determined (under these conditions inhibition is too high to accurately estimate a rate constant). Errors represent standard errors of the mean (s.e.m.) calculated from n = 3 independent experiments.

| [Intronistat B] ($\mu\text{M}$) | $k_1$ ($\text{min}^{-1}$) | $k_2$ ($\text{min}^{-1}$) | $k$ ($\text{min}^{-1}$) |
| --- | --- | --- | --- |
| 250 | N.D. | N.D. | N.D. |
| 100 | $0.003 \pm 0.001$ | N.D. | $0.003 \pm 0.001$ |
| 50 | $0.007 \pm 0.001$ | $0.001 \pm 0.000$ | $0.008 \pm 0.001$ |
| 20 | $0.004 \pm 0.001$ | $0.002 \pm 0.000$ | $0.006 \pm 0.001$ |
| 10 | $0.006 \pm 0.000$ | $0.002 \pm 0.000$ | $0.008 \pm 0.000$ |
| 1 | $0.022 \pm 0.004$ | $0.021 \pm 0.005$ | $0.043 \pm 0.007$ |
| 0.5 | $0.030 \pm 0.001$ | $0.025 \pm 0.001$ | $0.055 \pm 0.007$ |
| 0.05 | $0.042 \pm 0.004$ | $0.036 \pm 0.003$ | $0.078 \pm 0.007$ |
| 0 | $0.037 \pm 0.002$ | $0.031 \pm 0.003$ | $0.068 \pm 0.004$ |
| [di-Br-Intronistat B] ($\mu\text{M}$) | $k_1$ ($\text{min}^{-1}$) | $k_2$ ($\text{min}^{-1}$) | $k$ ($\text{min}^{-1}$) |
| 250 | $0.006 \pm 0.001$ | $0.006 \pm 0.001$ | $0.012 \pm 0.004$ |
| 100 | $0.012 \pm 0.001$ | $0.009 \pm 0.002$ | $0.021 \pm 0.003$ |
| 50 | $0.019 \pm 0.005$ | $0.015 \pm 0.005$ | $0.034 \pm 0.010$ |
| 20 | $0.020 \pm 0.004$ | $0.017 \pm 0.004$ | $0.037 \pm 0.006$ |
| 10 | $0.018 \pm 0.004$ | $0.013 \pm 0.004$ | $0.031 \pm 0.007$ |
| 1 | $0.028 \pm 0.004$ | $0.024 \pm 0.004$ | $0.052 \pm 0.008$ |
| 0.5 | $0.030 \pm 0.003$ | $0.023 \pm 0.003$ | $0.053 \pm 0.006$ |
| 0 | $0.037 \pm 0.002$ | $0.031 \pm 0.003$ | $0.068 \pm 0.004$ |

**Table S2:** X-ray data collection and refinement statistics (molecular replacement).

|  | OiD1-5 with<br>Mg<sup>2+</sup>, K<sup>+</sup> and<br>intronistat B | OiD1-5 with<br>Mg<sup>2+</sup>, K<sup>+</sup> and<br>di-Br-intronistat B | Oi5eD1-5 with<br>Ca<sup>2+</sup>, K<sup>+</sup> and<br>intronistat B | Oi5eD1-5 with<br>Mg<sup>2+</sup>, K<sup>+</sup> and<br>intronistat B | OiD1-5 with 5'-<br>exon, Mg<sup>2+</sup>, K<sup>+</sup><br>and intronistat B | OiD1-5 with<br>Mg<sup>2+</sup>, Na<sup>+</sup> and<br>intronistat B |
| --- | --- | --- | --- | --- | --- | --- |
| **PDB code** | 8OLS | 8OLV | 8OLW | 8OLY | 8OLZ | 8OM0 |
| **Data Collection** |  |  |  |  |  |  |
| Wavelength | 0.96 | 0.92 | 0.96 | 0.96 | 0.96 | 0.96 |
| Resolution<br>range<sup>a</sup> | 39.7 - 3.0<br>(3.1 - 3.0) | 48.4 - 2.8<br>(2.9 - 2.8) | 48.64 - 4.0<br>(4.1 - 4.0) | 48.02 - 3.1<br>(3.2 - 3.1) | 39.8 - 3.3<br>(3.4 - 3.3) | 42.3 - 3.6<br>(3.7 - 3.6) |
| Space group | *P* 2<sub>1</sub> 2<sub>1</sub> 2<sub>1</sub> | *P* 2<sub>1</sub> 2<sub>1</sub> 2<sub>1</sub> | *P* 2<sub>1</sub> 2<sub>1</sub> 2<sub>1</sub> | *P* 2<sub>1</sub> 2<sub>1</sub> 2<sub>1</sub> | *P* 2<sub>1</sub> 2<sub>1</sub> 2<sub>1</sub> | *P* 2<sub>1</sub> 2<sub>1</sub> 2<sub>1</sub> |
| Cell dimensions: |  |  |  |  |  |  |
| *a*, *b*, *c* (Å) | 89.0, 95.0, 223.9 | 90.0, 94.7, 225.0 | 88.1, 94.1, 222.9 | 88.5, 94.7, 222.9 | 87.7, 94.0, 223.4 | 88.8, 94.3, 223.9 |
| *α*, *β*, *γ* (°) | 90, 90, 90 | 90, 90, 90 | 90, 90, 90 | 90, 90, 90 | 90, 90, 90 | 90, 90, 90 |
| Total<br>reflections<sup>a</sup> | 207169<br>(20770) | 582731<br>(48555) | 74878<br>(21251) | 142551<br>(12794) | 183125<br>(18114) | 91376<br>(8942) |
| Unique<br>reflections<sup>a</sup> | 38577<br>(3799) | 46958<br>(4256) | 16065<br>(1442) | 34101<br>(3245) | 28270<br>(2786) | 21631<br>(2103) |
| Multiplicity<sup>a</sup> | 5.4<br>(5.5) | 12.4<br>(11.4) | 4.6<br>(4.8) | 4.2<br>(3.9) | 6.5<br>(6.5) | 4.2<br>(4.3) |
| Completeness<br>(%)<sup>a</sup> | 99.31<br>(99.48) | 97.90<br>(79.41) | 97.77<br>(85.51) | 98.69<br>(93.76) | 99.46<br>(98.71) | 96.82<br>(96.77) |
| Mean<br>I/sigma(I)<sup>a</sup> | 18.58<br>(0.92) | 11.97<br>(0.29) | 5.8<br>(0.9) | 10.57<br>(0.67) | 4.57<br>(0.47) | 12.17<br>(0.71) |
| **Refinement** |  |  |  |  |  |  |
| Wilson B-factor | 123.04 | 114.75 | 215.82 | 132.56 | 137.98 | 202.19 |
| R-merge<sup>a</sup> | 0.044<br>(2.0) | 0.11<br>(5.8) | 0.097<br>(2.1) | 0.069<br>(1.9) | 0.21<br>(2.7) | 0.043<br>(2.3) |
| R-meas<sup>a</sup> | 0.049<br>(2.2) | 0.115<br>(6.0) | 0.110<br>(2.4) | 0.080<br>(2.2) | 0.23<br>(3.0) | 0.049<br>(2.6) |
| R-pim<sup>a</sup> | 0.021<br>(0.89) | 0.034<br>(1.73) | 0.050<br>(1.08) | 0.040<br>(1.10) | 0.094<br>(1.16) | 0.023<br>(1.21) |
| CC<sub>1/2</sub><sup>a</sup> | 1.0<br>(0.35) | 1.0<br>(0.083) | 1.0<br>(0.28) | 1.0<br>(0.22) | 0.99<br>(0.32) | 1.0<br>(0.20) |
| No. reflections<sup>a</sup> | 38568<br>(3799) | 46400<br>(3718) | 15922<br>(1346) | 34019<br>(3187) | 28160<br>(2754) | 21611<br>(2099) |
| R-work<sup>a</sup> | 0.19<br>(0.43) | 0.21<br>(0.75) | 0.22<br>(0.40) | 0.20<br>(0.40) | 0.20<br>(0.41) | 0.23<br>(0.39) |
| R-free<sup>a,b</sup> | 0.25<br>(0.41) | 0.24<br>(0.69) | 0.25<br>(0.41) | 0.23<br>(0.37) | 0.27<br>(0.42) | 0.28<br>(0.40) |
| No. atoms: |  |  |  |  |  |  |
| RNA | 8349 | 8438 | 8497 | 8475 | 8455 | 8349 |
| ligand/ion | 145 | 115 | 21 | 107 | 112 | 45 |
| water | 13 | 11 | 11 | 49 | 15 | 0 |
| B-factor: |  |  |  |  |  |  |
| RNA | 144 | 126 | 222 | 146 | 146 | 226 |
| ligand/ion | 174 | 160 | 208 | 125 | 143 | 199 |
| water | 104 | 91 | 168 | 119 | 82 |  |
| RMSD: |  |  |  |  |  |  |
| bond length (Å) | 0.042 | 0.018 | 0.011 | 0.011 | 0.030 | 0.042 |
| bond angles (°) | 2.15 | 1.89 | 2.22 | 1.69 | 1.90 | 2.65 |

One single crystal was used to collect each data set, respectively.
<sup>a</sup>Values in parentheses are for the highest-resolution shell.
<sup>b</sup>R-free was calculated using 5% of the total reflections.

### Supplemental Scheme

**Scheme S1.**
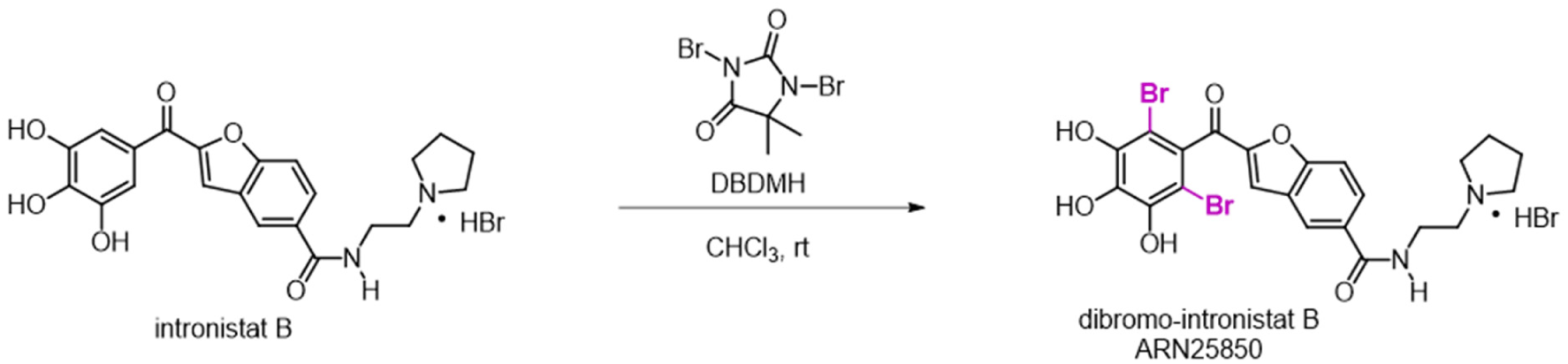
Chemical synthesis of dibromo-intronistat B hydrobromide, ARN25850.

